# Transcription controls chromatin-nuclear lamina contacts through distinct Lamin A and LBR tethering mechanisms

**DOI:** 10.64898/2026.07.14.738400

**Authors:** Marco Bernasconi, Jeremie Breda, Tom van Schaik, Anna G Manjon, Federico Zambelli, Giulio Pavesi, René H Medema, Marco Muzi Falconi, Bas van Steensel, Stefano G Manzo

## Abstract

Lamina-associated domains (LADs) are large genomic regions that interact with the nuclear lamina (NL). Much of the underlying “grammar” governing their positioning at the nuclear periphery remains unclear. LADs are composed of heterochromatin and typically harbor repressed genes, and their association with the NL is generally incompatible with strong transcriptional activity. The extent to which transcription globally shapes chromatin-NL interactions is not fully understood. Here, we combined acute transcription inhibition using Flavopiridol or Triptolide with genome-wide mapping of chromatin-NL contacts. We found that chromatin-NL interactions are rapidly rewired upon transcription inhibition. Changes in chromatin-NL contacts upon transcription shutdown are predictable based on transcriptional activity and the presence of H3K9me3-marked heterochromatin. This rewiring is reversible, as genome-NL interactions quickly return to baseline levels following drug wash-off. Notably, gain and loss of chromatin-NL interactions upon transcription shutdown reflect two distinct tethering mechanisms. Inter-LADs genomic regions (iLADs) enriched in highly active genes and located near stable LADs, which are tethered by Lamin A (LMNA/C), become re-attached to the NL following transcription inhibition. In parallel, H3K9-methylated regions tethered to the nuclear envelope by the Lamin B receptor (LBR) undergo extensive detachment from the NL. Strikingly, LMNA/C and LBR oppositely regulate transcription-sensitive LADs and are required for transcriptional control of chromatin–NL contacts. Together, our findings highlight the plasticity and dynamic nature of chromatin-NL interactions and provide the first evidence that LMNA/C- and LBR-mediated tethering mechanisms exhibit distinct sensitivities to transcription inhibition.

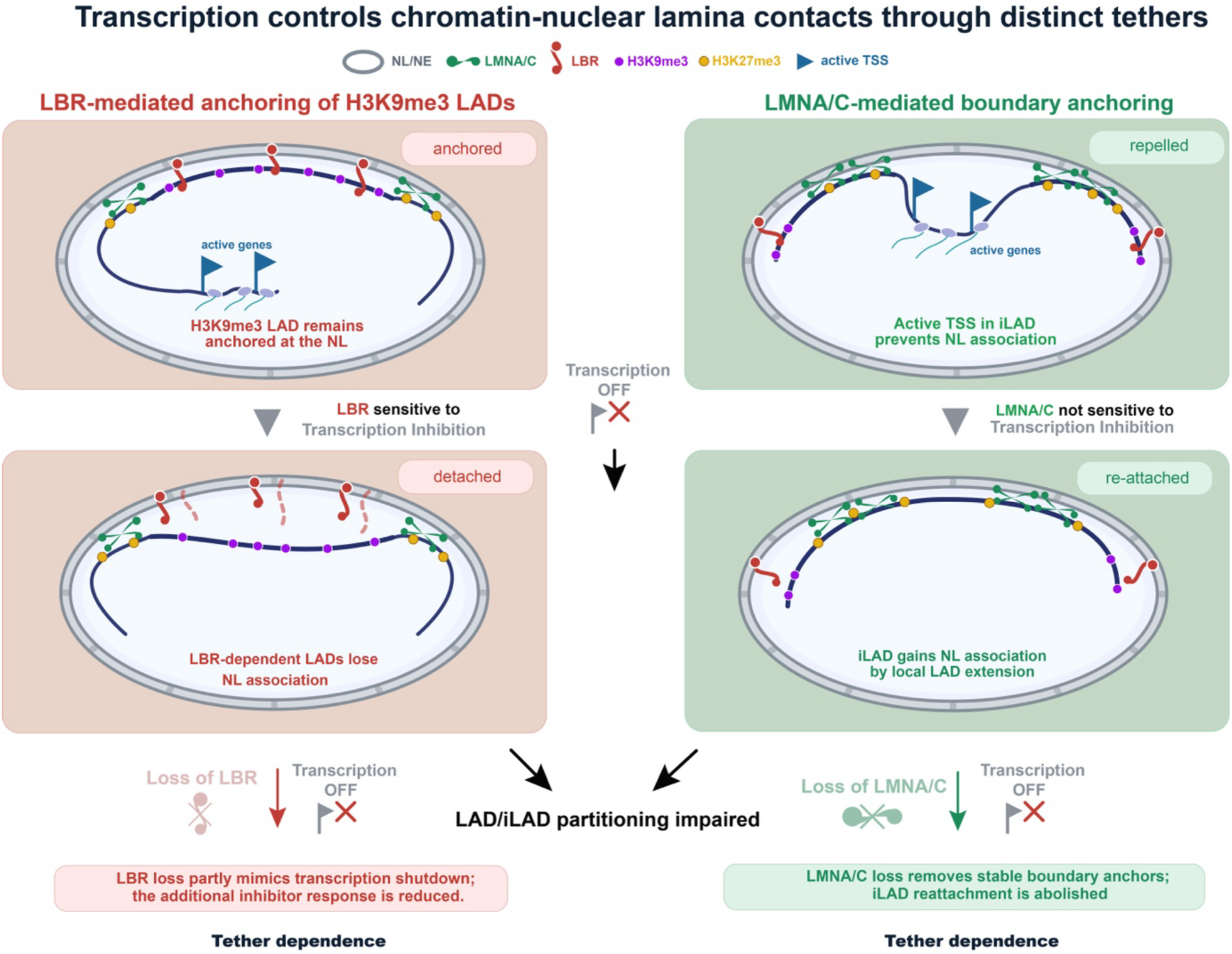

**HIGHLIGHTS:**

- Transcription inhibition alters chromatin–NL contacts rapidly and reversibly
- Active transcription prevents inter-LADs located near LMNA/C-tethered LADs from associating with the nuclear lamina.
- LBR-tethered heterochromatin is repositioned away from the NL
- Transcription-dependent modulation of chromatin-NL contacts is dependent on LMNA/C and partially on LBR

## INTRODUCTION

The spatial organization of the genome within the nucleus is tightly linked to gene regulation. A prominent feature of nuclear architecture is the association of large genomic regions with the nuclear lamina (NL), forming lamina-associated domains (LADs). LADs are typically gene-poor, enriched in repressive chromatin marks, largely transcriptionally inactive, and have therefore been proposed to represent a repressive nuclear compartment ^1–4^. While some LADs are constitutively associated with the NL across cell types, others display more dynamic, cell- type-specific interactions, indicating that genome-NL contacts are regulated and context-dependent ^5–7^. However, the mechanisms governing this regulation remain poorly understood.

Multiple lines of evidence support a functional link between lamina association and transcriptional repression. During differentiation, reshaping of chromatin-NL interactions is coordinated with changes in the transcriptional program ^8–11^. These observations support a reciprocal relationship between transcription and genome positioning.

Several studies suggest that LADs constitute a strong repressive environment for transcription. Simultaneous measurement of NL association and gene expression in single cells revealed that contact frequency with the NL inversely correlates with gene expression^12^. Multiple parallel reporter assays show that reporters integrated into LADs have lower transcriptional output than those integrated into inter-LAD (iLADs) locations ^13^. Similarly, when promoters of LAD genes are transferred into a non-LAD context, they become activated ^14^. Artificial tethering of genomic loci to the NL can reduce gene expression in some ^4,15^, but not all cases ^16^. Finally, relocation of genes away from LADs or lamina depletion correlate with transcriptional activation ^17,18^.

Transcription seems to be a state that is not compatible with physical tethering to the NL. Gene activation within LADs has been shown to trigger local detachment from the NL, suggesting that transcription may antagonize lamina association ^19–21^

Importantly, most evidence linking transcription and NL interactions derives from locus-specific perturbations or steady-state comparisons. Although very informative, these studies do not address how acute and global changes in transcription affect chromatin-lamina interactions across the genome.

The use of transcription inhibitors combined with high-throughput approaches has further highlighted the functional connection between NL association and transcription. Inhibition of RNA Polymerase II, using alpha-amanitin increases association with the NL for a few selected loci ^22^. In early developing embryos, altering zygotic genome activation with transcription inhibitors affects LAD organization ^23^. Finally, inhibiting transcription elongation with DRB affects a subset of small LADs that depend on the architectural factor CTCF ^24^. All these studies have provided valuable insights into how transcription inhibition might affect chromatin-NL contacts. However, both the use of alpha-amanitin and DRB come with limitations. Alpha-amanitin has notoriously low uptake and requires long-term treatment ^25^, not allowing for capturing the immediate effect of transcription inhibition. DRB is faster but not potent or specific ^25,26^. It is likely that immediate or stronger effects might have been missed, and the use of more potent and fast-acting transcription inhibitors ^25,26^ might reveal additional information on how LADs and iLADs dynamically respond to rapid and potent transcriptional shutdown. Despite many efforts, a study systematically dissecting how transcription directly shapes chromatin-NL interactions remains missing.

An additional layer of complexity arises from the presence of multiple lamina-tethering mechanisms. Peripheral heterochromatin can be anchored through several tethering mechanisms, including the Lamin B receptor (LBR) and Lamin A/C (LMNA/C). These two tethers have been proposed to act through distinct or partially redundant pathways ^27,28^. While these tethering systems are known to play key roles during development, their respective contributions to dynamic genome (re)organization in relation to gene transcription is unclear.

Here, we address these questions by combining acute transcription inhibition with genome-wide mapping of chromatin-NL interactions. Using pA-DamID ^29^ after rapid transcription perturbation, we uncover a strikingly dynamic and reversible reorganization of genome-NL contacts. We show that transcription inhibition induces both gain and loss of lamina association across the genome, revealing a striking plasticity in LAD organization. Importantly, these changes are not uniform but instead reflect two opposing and mechanistically distinct tethering pathways. Transcriptionally active regions preferentially gain lamina association in proximity of LMNA/C-dependent regions, whereas H3K9me3-enriched heterochromatin loses lamina association in a manner consistent with LBR-dependent tethering.

Together, our findings establish transcription as a central regulator of genome organization at the nuclear lamina and reveal that its effects are mediated through distinct tethering pathways with opposing responses to transcriptional perturbation. These results provide a conceptual model for understanding how genome-lamina interactions are dynamically maintained through the interplay between transcriptional activity and alternative chromatin tethering mechanisms.

## RESULTS

### Transcription controls genome partitioning in LADs and iLADs

#### Transcription inhibitors affect chromatin-NL contact

To investigate how transcription shapes chromatin–NL interactions, we combined short-term treatments with transcription inhibitors and pA-DamID, a method that enables mapping of chromatin–NL contacts following rapid perturbations ^29^. We used two potent transcription inhibitors: flavopiridol (FLV), which inhibits CDK9 and impairs transcription elongation ^26^, and Triptolide (TPL), which binds the TFIIH subunit XPB and triggers CDK7-dependent degradation of RNA Polymerase II ^30,31^. Cells were treated for three hours (**Supplementary Figure 1A**), followed by pA-DamID using an antibody against LMNB2, a B-type lamin component of the nuclear lamina.

Both TPL and FLV induced widespread gain and loss of chromatin–NL interactions (**Figure 1A**). Notably, both treatments caused a pronounced shift in the distribution of the LMNB2 signal, from the near-bimodal profile observed in control cells to a predominantly unimodal distribution (**Figure 1B**). Upon transcription inhibition, LADs exhibited a significant reduction in NL interaction scores, whereas iLADs showed a marked increase in NL association, although the effect was variable across different regions (**Figure 1C**). These data indicate that transcription inhibition disrupts genome partitioning between these two compartments. Analysis at the LAD boundaries showed that LAD borders are not massively repositioned, as a clear demarcation between iLADs and LADs remains visible after inhibition, although weaker (**Figure 1D**).

**Figure 1.**
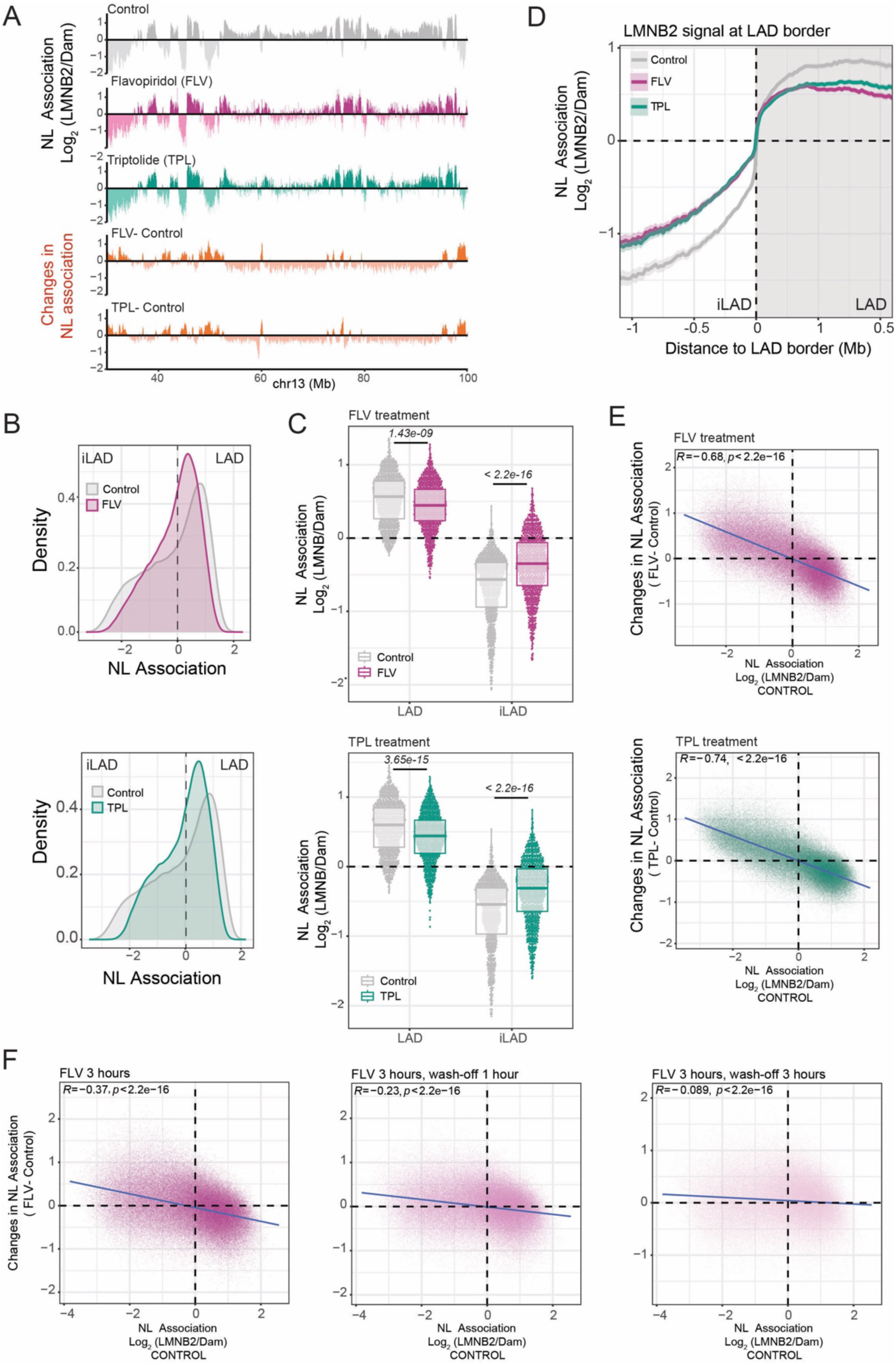
Transcription controls genome partitioning in LADs and iLADs. A) *Top:* Representative genomic track of LMNB2 pA-DamID for control sample (grey), flavopiridol (purple), triptolide (green), treatments. 20-kb bins were used. The antibody signal is normalized over a Dam-only control. *Bottom:* Differential tracks (knockdown - control, orange) highlighting the gain and loss of genome- NL interactions following inhibition of transcription with the two different drugs. B) LMNB2 signal distribution for control and flavopiridol (top panel) and triptolide (bottom panel). In control cells, a bimodal distribution usually represents iLAD/LAD partitioning. Results are from 4 independent biological replicates for FLV and 3 for TPL. C) LAD and iLAD score following FLV or TPL treatment using LMNB2 mapping data. Results are from 4 biological replicates. D) Average LMNB2 pA-DamID score around LAD borders for control and FLV or TPL treated samples. Genome-NL contact profiles are the average of 4 biological replicates. The solid line and the shaded area represent the mean signal and 95% confidence interval of the mean, respectively. E) Correlation scatter plots of 20 kb genomic bins for LMNB2 score in control cells (x-axis) and differential LMNB2 score (transcription inhibitor - control, y-axis) for FLV and TPL treatments. The blue line represents a linear model; Pearson correlation, and *p*-value are shown in the plots. F) Same plot but for the wash-off experiment where cells were treated for 3 hours with FLV (left panel) and then drug was removed, and chromatin-NL interactions were probed at 1 hour (middle panel) and 3 hours (right panel) post drug removal. Results are from 2 biological replicates. For all experiments, drug treatment was performed at 3 hours and 2 μM.

The de-partitioning effect was further quantified by examining the slope of the correlation between NL association in control cells and changes induced by transcription inhibition (**Figure 1E**). Both TPL and FLV produced highly similar effects, as reflected by the strong correlation between their induced changes (**Supplementary Figure 1B**). Consistent results were obtained when mapping chromatin interactions with LMNB1, indicating that these effects reflect changes at the level of the nuclear lamina rather than a specific lamina component (**Supplementary Figure 1C**). Finally, calibrated pA-DamID experiments using spiked-in mouse cells yielded results nearly identical to uncalibrated data, ruling out normalization biases (**Supplementary Figure 1D**).

Together, these findings demonstrate that transcription plays a key role in maintaining genome partitioning between the nuclear interior and the nuclear lamina.

### Effects of transcription inhibition on chromatin-NL contacts are rapid and reversible

#### Transcription inhibition rapidly affects chromatin–NL contacts

To assess how dynamically these interactions respond to transcriptional perturbation, we performed a time-course experiment using both TPL and FLV. Notably, we found that most of the de-partitioning effect was already evident after one hour of transcription inhibition (**Supplementary Figure 1E**), with both LADs and iLADs reaching NL interaction scores comparable to those observed at the three-hour time point (**Supplementary Figure 1F**). These results indicate that transcription inhibition rapidly alters chromatin–NL contacts and support a direct role for transcription in maintaining genome partitioning between the nuclear lamina and the nuclear interior.

#### The effects of transcription inhibition are reversible

To investigate the reversibility of these effects, we performed wash-off experiments using FLV, whose binding to CDK9 and inhibitory effects on transcription are reversible. Cells were treated with FLV for three hours, followed by drug removal, and chromatin–NL contacts were measured at multiple time points thereafter. We observed a progressive restoration of chromatin–NL interactions, with most contacts returning to control levels within three hours after wash-off (**Figure 1F** and **Supplementary Figure 1G**). Together, these findings demonstrate that changes in chromatin–NL contacts induced by transcription shutdown are highly dynamic and reversible, highlighting the plasticity of genome organization in response to transcriptional perturbation.

### Chromatin composition is predictive of chromatin-NL interaction changes

#### Gain and loss of chromatin–NL contacts reflect distinct chromatin compositions

LADs are repressive regions, but distribution of heterochromatic markers in LADs is not random ^3,32,33^. To determine whether transcription inhibition perturbs chromatin–NL interactions in a chromatin feature–specific manner, we generated pA-DamID maps for three major repressive histone marks: H3K9me3, H3K9me2, and H3K27me3. These datasets were complemented with transcriptomic data previously generated in RPE1 cells ^32^. We integrated these four datasets using principal component analysis (PCA) to assess the predictive contribution of each feature to genome-wide chromatin variability (**see method section**).

PCA revealed that the first two principal components account for 92% of the total variance, with PC1 explaining 56% and PC2 explaining 36%. This indicates that most of the variability across these datasets can be captured by two dominant and largely independent axes, reflecting coordinated patterns between gene expression and repressive histone modifications (**Figure 2A**, and **Supplementary Figure 2A**). Overlaying these results with differential LMNB2 signals between transcription inhibitor–treated (FLV or TPL) and control cells revealed distinct chromatin-specific responses. Transcriptionally active regions preferentially gained chromatin–NL interactions upon transcription inhibition, consistent with previous observations ^21,22,34^. In contrast, H3K9me3-enriched heterochromatin predominantly lost chromatin–NL interactions, whereas regions marked by H3K9me2 or H3K27me3 did not show a consistent trend (**Figure 2A**). This behavior of H3K9me3-marked heterochromatin was further confirmed by direct correlation analysis (**Supplementary Figure 2B**). Together, these results indicate that transcription inhibition induces two major chromatin reorganization events: reattachment of transcriptionally active regions to the NL and displacement of H3K9me3-enriched heterochromatin away from the NL.

**Figure 2.**
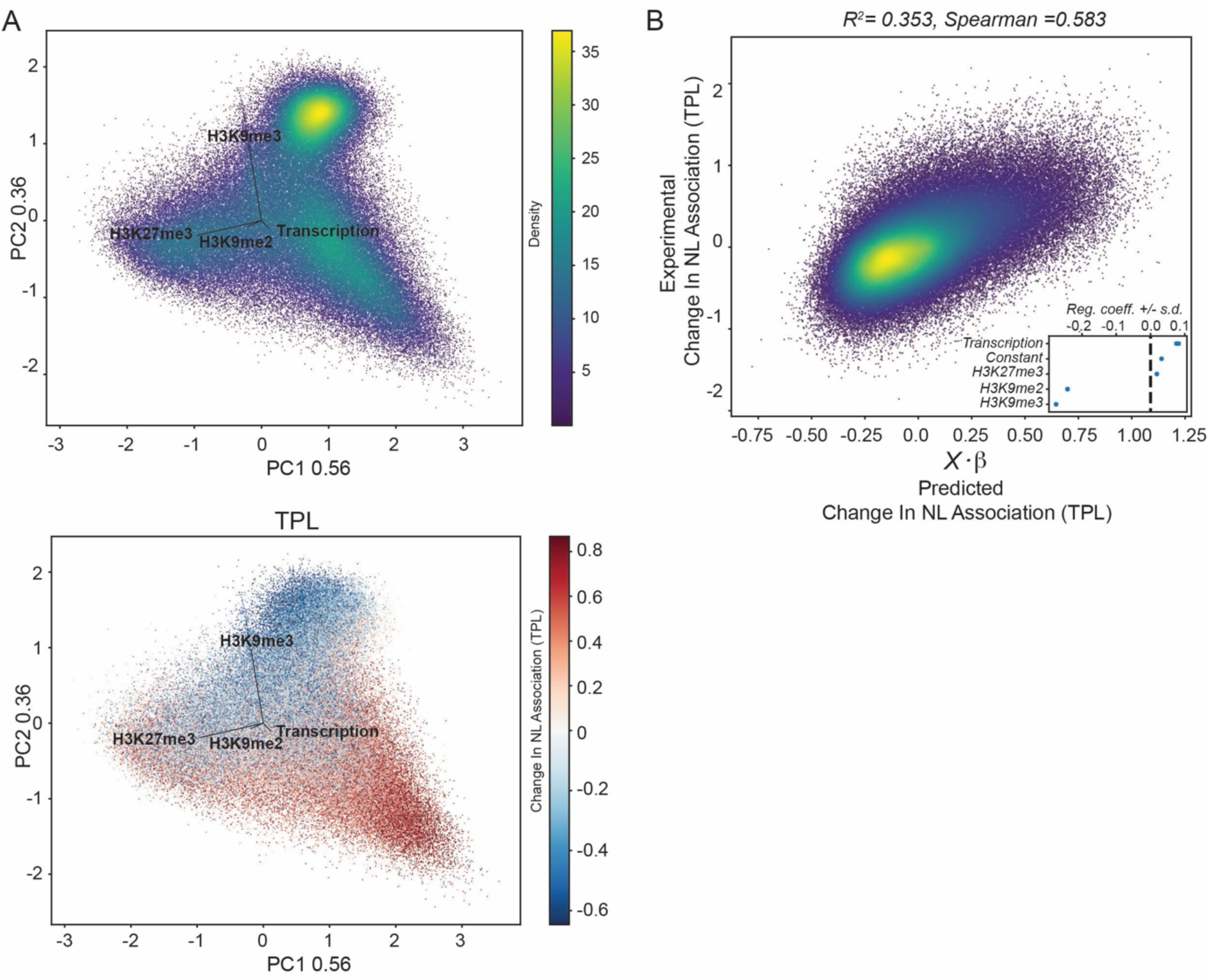
Chromatin composition predicts changes in chromatin–nuclear lamina interactions. **A)** Top panel: PCA performed on 100-kb genomic bins using H3K9me2, H3K9me3 and H3K27me3 pA-DamID data together with RNA-seq data from RPE1 cells. H3K9me2, H3K9me3, H3K27me3 and log-transformed RNA-seq values were decomposed by singular value decomposition. PC1 and PC2 are shown and together explain 92% of the variability in chromatin composition. Dots are colored by density, and black arrows indicate the feature loadings for each marker. *Bottom panel:* Same PCA analysis as in A, with bins colored by changes in chromatin–NL interactions following TPL treatment relative to control conditions (TPL–Control; 3 h, 2 μM). **B)** Least-squares regression model of TPL-induced changes in chromatin–NL interactions. Observed changes in nuclear lamina association are plotted against least-squares predicted values, calculated as X.β from a linear model including an intercept plus H3K9me2, H3K9me3, H3K27me3 and RNA-seq. The model shows that transcription and H3K9me3-marked heterochromatin predict gain and loss of chromatin–NL interactions following TPL treatment, respectively. R² and Spearman correlation are shown on the plot. The inset reports the estimated regression coefficients for each predictor, with ±2 standard-deviation intervals. Results are from two biological replicates.

#### Transcriptional activity and heterochromatin features predict chromatin–NL dynamics

To quantitatively assess the predictive power of individual chromatin features, we applied least- squares linear regression modeling. Despite minor differences between treatments, both TPL and FLV yielded consistent results: transcriptional activity emerged as a positive predictor of NL association upon transcription inhibition, whereas H3K9me3 enrichment, (and to a minor extent, H3K9me2), were negative predictors (**Figure 2B** and **Supplementary Figure 2C**). Model performance was supported by high Spearman correlation coefficients for both treatments (**Figure 2B** and **Supplementary Figure 2C**), indicating robust predictive power.

Together, these findings demonstrate that transcriptional output and H3K9me3 deposition are key predictors of how genomic regions reorganize their association with the nuclear lamina following transcriptional shutdown.

### Transcription mantains highly transcribed iLADs away from NL

*A significant fraction of iLADs gain chromatin–NL interactions upon transcription inhibition*. Although increased chromatin–NL interactions originate from active genes (**Supplementary Figure 3A**), this gain extends beyond individual gene units and can encompass entire iLADs (**Figure 1A**). To quantify the number of regions affected globally, we performed Limma-Voom differential analysis to identify iLADs with significant changes in NL association ^29,32^. Of the 889 iLADs identified in RPE1 cells, 358 (∼40%) showed a significant increase in NL interactions following TPL treatment (**Figure 3A**). Similarly, 299 iLADs (∼34%) displayed increased NL association upon FLV treatment (**Supplementary Figure 3B**). In contrast, only a small fraction of iLADs exhibited further decreases in NL interactions (45 iLADs, ∼5% for TPL; 18 iLADs, ∼2% for FLV) (**Figure 3A** and **Supplementary Figure 3B-C**). These results indicate that a substantial portion of inter-LAD regions increases its association with the nuclear lamina upon transcription inhibition.

**Figure 3.**
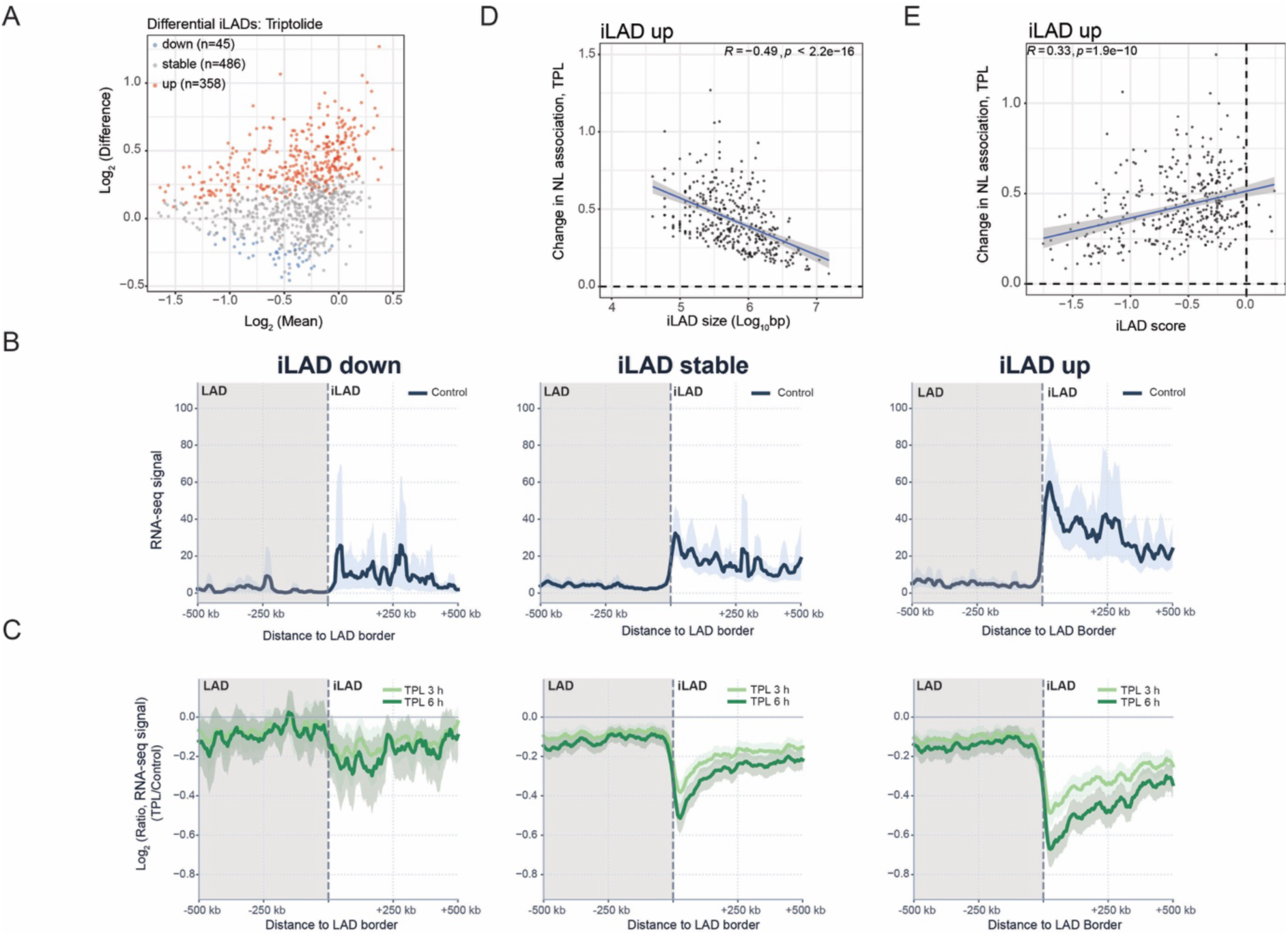
Highly active genes proximal to LAD borders associate with the NL after transcription inhibition. A) Results of Voom- Limma analysis used to call iLADs that significantly gain or lose interaction with LMNB2 following TPL treatment. Results are from 4 biological replicates. B) Average calibrated RNA-seq signal (binned bam coverage) around LAD/iLAD borders in control cells for the three different iLAD categories (stable, down, and up) identified in Figure 3A. RNA-seq profiles are the average of 3 biological replicates. The solid line and the shaded area represent the mean signal and 95% confidence interval of the mean, respectively. C) Same as B, but for log2 (ratio) of RNA-seq signal between TPL treatment (3 and 6 hours, 2 μM) and control. D) Correlation between *gained* iLAD size (Log10(bp)) and differential LMNB2 score (transcription inhibitor - control, y-axis) for TPL treatment. E) Correlation between *gained* iLAD LMNB2 pA-DamID score in control cells and differential LMNB2 score (transcription inhibitor - control, y-axis) for TPL treatment. For D and E, the blue line represents a linear model; the Pearson correlation and p-value are shown in the plots.

To directly link changes in chromatin–NL contacts to transcriptional output, we performed calibrated RNA-seq. RPE1 cells were treated or not with TPL for 3 and 6 hours and spiked with a fixed percentage of Drosophila cells (**Supplementary Figure 3D–E**). Calibrated data revealed a pronounced global downregulation of transcripts at both time points, an effect that could not be accurately captured without spike-in normalization (**Supplementary Figure 3F–G**).

Aggregated analysis of RNA-seq signal distribution at LAD/iLAD borders for the three different iLAD categories revealed that iLADs showing increased interaction following transcription inhibition harbor a higher level of transcription compared to stable iLADs or iLADs down (iLAD up). This was particularly true near LAD borders (**Figure 3B**).

Similar results were obtained by analyzing genes: we computed an average genic LMNB2 score for control and treated cells and classified genes into three categories: gain (ΔLMNB2 score inhibitor–control > 0.5), loss (ΔLMNB2 score < −0.5), and no change (−0.5 < ΔLMNB2 score < 0.5). This approach identified few thousand genes exhibiting descrete gains or losses in chromatin–NL contacts (**Supplementary Figure 3H**). Analysis of gene expression across these categories revealed that genes gaining NL interactions are highly expressed, with significantly higher expression levels than stable genes or those showing reduced chromatin– NL interactions upon treatment (**Supplementary Figure 3I**).

Intriguingly, when comparing transcriptional changes induced by TPL across iLAD categories, the strongest downregulation was observed in regions that gained NL interactions (**Figure 3C** and **Supplementary Figure 3L**) especially in proximity of LAD borders. We did not detect a strong global correlation between transcriptional decrease and changes in chromatin–NL contacts for genes (**Supplementary Figure 3M**).

Together, these results indicate that genes repositioned to the NL following transcription shutdown are those most affected in terms of transcriptional output.

#### Small iLADs proximal to stable LADs exhibit a greater propensity for NL association

We next investigated which genomic and chromatin features characterize iLADs that gain NL interactions, aiming to identify determinants of their sensitivity to transcription inhibition. We hypothesized that smaller iLADs, which by definition are flanked by closely spaced LAD borders, may be more prone to re-association with the NL. Consistent with this idea, we observed a clear inverse correlation between iLAD size and the magnitude of NL reattachment, with smaller iLADs showing stronger increases in NL association (**Figure 3D**). Similar results were obtained by an analysis performed on genes where we detected a modest but significant correlation between proximity to LAD borders and increased chromatin– NL interactions, such that genes located closer to LAD boundaries show greater gains in NL association (**Supplementary Figure 4A-D**).

We further examined whether the baseline NL interaction propensity of iLADs influences their response. Indeed, we found a positive correlation between the increase in NL association and the LMNB2 interaction score in control cells, indicating that iLADs with higher basal NL affinity are more likely to reattach upon transcription inhibition (**Figure 3E**).

#### iLADs reattached to the NL do not undergo heterochromatinization

We next asked whether iLADs that reposition to the NL upon transcription inhibition undergo changes in chromatin state characteristic of LADs. To address this, we mapped the distribution of the major heterochromatin marks (H3K9me3, H3K9me2, and H3K27me3) in control and TPL-treated cells. Despite their repositioning toward the NL, gained iLADs did not show substantial changes in these chromatin features (**Supplementary Figure 4E**). Thus, at the analyzed time points, iLADs that re-associate with the NL do not undergo heterochromatinization.

Although we cannot rule out that heterochromatinization might occur at longer timepoints following transcription inhibition, these results support a model in which transcription primarily prevents NL association of small iLADs exhibiting a higher intrinsic propensity to interact with the nuclear lamina, but that do not undergo heterochromatinization.

### iLADs protected by transcription are proximal to LMNA/C-tethered regions

#### LAD borders adjacent to gained iLADs are stable

To further explore the role of genomic context, we analyzed the LMNB2 profiles of iLADs and their surrounding LAD regions across the three iLAD categories defined above. iLADs gaining NL interactions showed a strong increase in LMNB2 signal compared to iLADs stable or iLADs down (**Figure 4A**, and **Supplementary Figure 4F-G**). Notably, LADs flanking gained iLADs displayed relatively stable NL interaction profiles, whereas LADs adjacent to stable or weakened iLADs exhibited slightly reduced NL interactions (**Figure 4A**, and **Supplementary Figure 4H**). These observations suggest that reattachment of inter-LAD regions preferentially occurs near stably anchored LADs.

**Figure 4.**
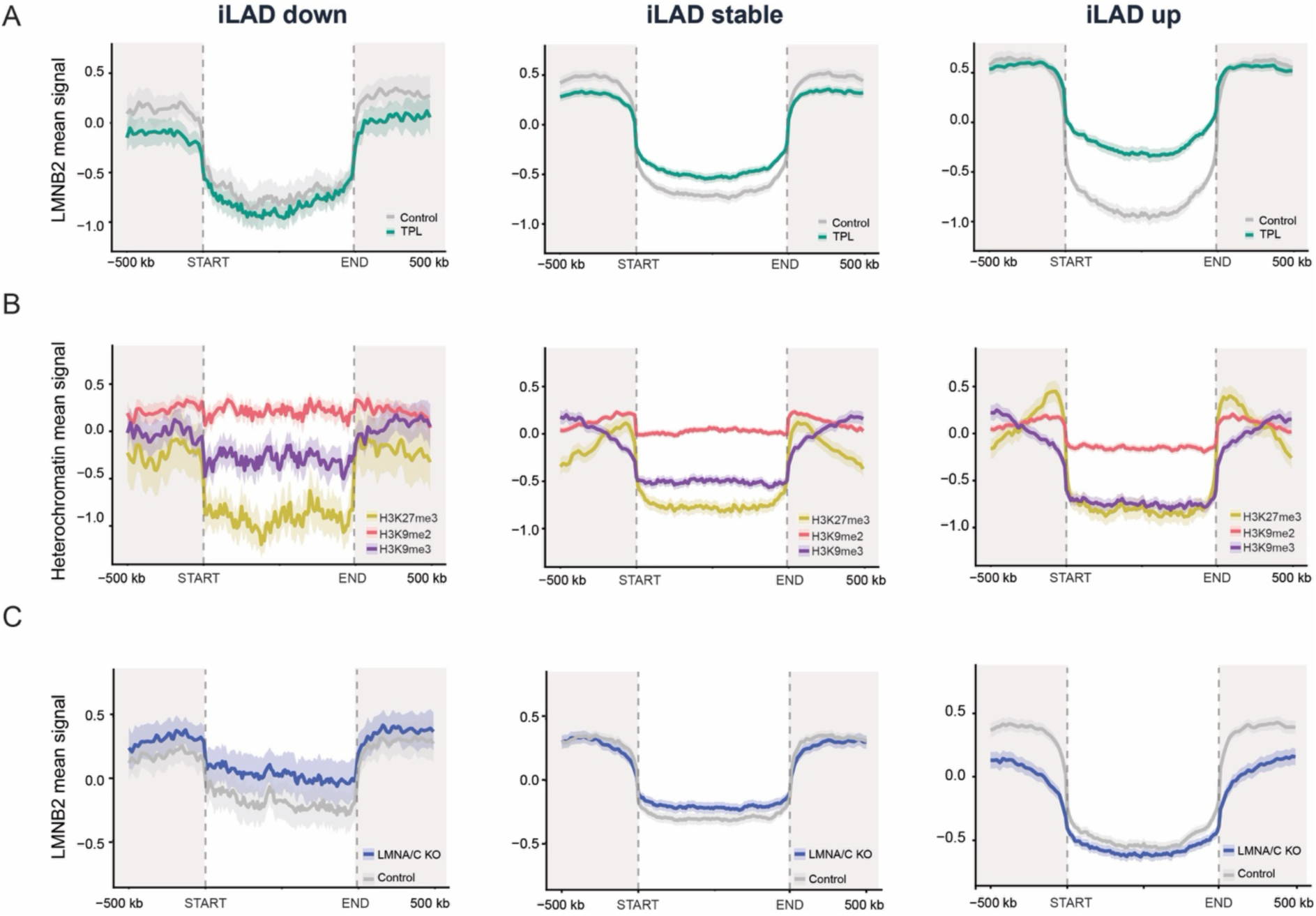
Transcription protects iLADs proximal to LMNA/C-thethered regions from association with the NL. A) Metaplot showing average pA-DamID signal for LMNB2 at iLADs up (*left*), stable (*middle*), or down (*right*), identified in Figure 3A, in control or TPL-treated cells (3 hours, 2 μM). Control is shown in grey, and TPL-treated samples are shown in green. iLADs were length-normalized, and the average antibody signal was computed. The start and end of the normalized iLAD are shown by dashed lines. 500 Kb of not-normalized LAD regions upstream and downstream of iLADs are shown in grey. The shaded area around the average antibody signal represents 95% confidence intervals. Results are from 4 biological replicates in RPE1 cells. B) Similar metaplots showing average pA-DamID signal in control cells for the three main heterochromatin components (H3K9me2, H3K9me3, H3K27me3) at the 3 iLADs categories identified in Figure 4A. Results are from 2 biological replicates. C) Same as A, but for Control (grey) and LMNA KO cells (blue). Results are from 4 biological replicates.

#### Stable LAD borders adjacent to gained iLADs are associated with LMNA/C-dependent tethering

To identify features distinguishing iLADs that reattach to the NL, we analyzed heterochromatin profiles across iLAD categories and their flanking LAD regions. As expected, iLADs were depleted of heterochromatic marks, especially H3K9me3 and H3K27me3, which increased sharply upon entry into LAD regions (**Figure 4B**).

Notably, LAD borders adjacent to gained iLADs were enriched in H3K27me3, a feature not observed at borders flanking stable or weakened iLADs (**Figure 4B**). H3K27me3 enrichment at LAD borders has been linked to LMNA/C-mediated tethering ^35–37^. We therefore hypothesized that the stability of these regions depends mainly on LMNA/C. To test this, we performed LMNB2 pA-DamID in LMNA/C knockout cells ^38^ (**Supplementary Figure 4G**). Strikingly, LADs adjacent to gained iLADs were sensitive to LMNA/C loss, indicating that LMNA/C is responsible, at least partially, for tethering this specific subset of regions. In contrast, LADs flanking stable or weakened iLADs were largely unaffected (**Figure 4C**).

Together, these results indicate that transcription repels inter-LAD regions located near LADs which are partially dependent on LMNA/C-mediated anchoring.

### LADs weakening without heterochromatin disruption

#### A subset of LADs weakens their interaction with the nuclear lamina without heterochromatin disruption

A prominent feature of the rapid rewiring of genome–NL interactions upon transcription inhibition is the weakening of a substantial fraction of LADs (**Figure 1B–D**), in agreement with previous results ^24^. This effect was statistically significant for 182 out of 900 LADs following TPL treatment and for 137 LADs upon FLV treatment, as determined by Limma-Voom analysis (**Figure 5A, Supplementary Figure 5A-B**). Analysis of H3K9me3, H3K9me2, and H3K27me3 levels in control and TPL-treated cells revealed that, despite a marked reduction in chromatin-NL contacts, weakened LADs exhibit minimal changes in heterochromatin marks (**Supplementary Figure 5C**). This suggests that, upon transcription inhibition, LAD-associated heterochromatin is repositioned away from the NL without undergoing major heterochromatin remodeling. However, we detected a modest positive correlation between changes in chromatin–NL interactions and variations in heterochromatic marks, particularly at LAD regions (**Supplementary Figure 5D**), indicating that subtle chromatin changes may accompany spatial repositioning.

**Figure 5.**
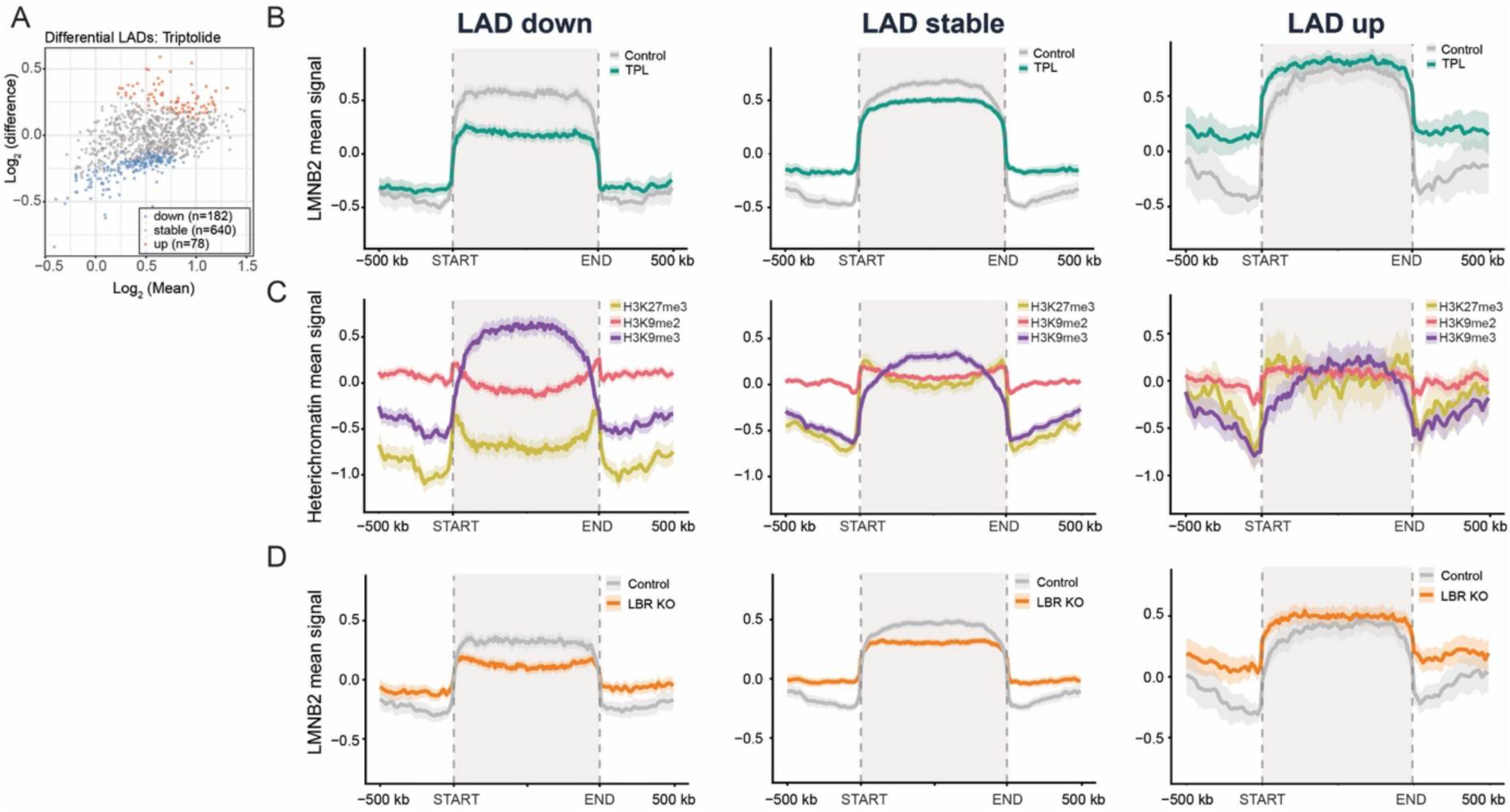
LBR tethers LADs reinforced by transcription. A) Results of Voom-Limma analysis used to call LADs that significantly gain or lose interaction with LMNB2 following TPL treatment (3 hours, 2 μM). Results are from 4 biological replicates. B) Metaplot showing average pA-DamID signal for LMNB2 at LADs down (*left*), stable (*middle*), or up (*right*) identified in Figure 4A, in control or TPL-treated cells (3 hours, 2 μM). Control is shown in grey, and TPL-treated samples are shown in green. LADs were length-normalized, and the average antibody signal was computed. The start and end of the normalized LADs are shown by dashed lines, and LAD regions are colored in grey. 500 Kb of not-normalized iLAD regions upstream and downstream of iLADs are shown in white. The shaded area around the average antibody signal represents 95% confidence intervals. Results are from 4 biological replicates in RPE1 cells. C) Similar metaplots showing average pA-DamID signal in control cells for the three main heterochromatin components (H3K9me2, H3K9me3, H3K27me3) at the 3 LADs categories identified in Figure 5A. Results are from 2 biological replicates. D) Same as B and C, but for Control (grey) and LBR KO cells (orange). Results are from 2 biological replicates.

#### Weakened LADs do not reflect local LAD expansion

We next asked whether the detachment of LADs upon transcription inhibition is associated with local expansion or contraction of LADs. To address this, we performed a meta-analysis analogous to that used for iLADs, comparing gained, stable, and weakened LADs.

This analysis revealed that weakened LADs do not exhibit coordinated extension of LAD borders, and gain of interactions in iLADs occurs mainly at stable LADs or LADs that slightly increase the interaction with the NL (**Figure 5B** and **Supplementary Figure 5E-F**).

These findings indicate that the gain of chromatin-NL interactions in iLADs and the loss of interactions in LADs occur through independent mechanisms rather than coordinated remodeling within the same genomic regions.

### LBR tethers LADs reinforced by transcription

#### LADs weakened by transcription inhibition are also affected by knock-out of LBR

We hypothesized that LADs undergoing detachment upon transcription inhibition may rely on a specific tethering mechanism sensitive to transcriptional activity. Analysis of chromatin features across LAD categories revealed that regions losing NL interactions are strongly enriched in H3K9me3 (**Figure 5C**), consistent with our PCA results. Given that H3K9me3- marked chromatin can be tethered by the Lamin B receptor (LBR) ^39^, we tested whether LADs down depend on LBR. Using our previously generated pA-DamID data in LBR knockout (LBR KO) cells ^32^, we compared LMNB2 profiles between wild-type and LBR KO cells across LAD categories. Strikingly, LADs that lose NL interactions upon transcription inhibition showed the strongest detachment in LBR KO cells, indicating a marked dependence on LBR-mediated tethering (**Figure 5D**).

Interestingly, both LBR loss and transcription inhibition also induced increased NL association in a subset of LADs (**Figure 5B** and **Figure 5D**), suggesting a functional link between LBR-mediated tethering and transcriptional activity.

### Opposite role of LBR and LMNA/C on LAD/iLAD partitioning

#### LMNA/C and LBR exert opposing effects on transcription-sensitive LADs

We next directly compared the effects of LMNA/C and LBR loss on transcription-sensitive LADs. LADs that detach upon transcription inhibition and depend on LBR exhibit increased NL interactions upon LMNA/C knockout (**Figure 6A**). This suggests that these regions are oppositely regulated by LMNA/C and LBR. Conversely, LADs that gain NL interactions following transcription inhibition show strong dependence on LMNA/C and increased interaction following LBR knock-out (**Figure 6A**). Moreover, genomic regions surrounding these LADs display increased NL interactions within adjacent iLADs, consistent with the idea that iLAD reattachment induced by transcription inhibition originates from LMNA/C-tethered regions (compare **Figure 4C** and **Figure 6A**).

**Figure 6.**
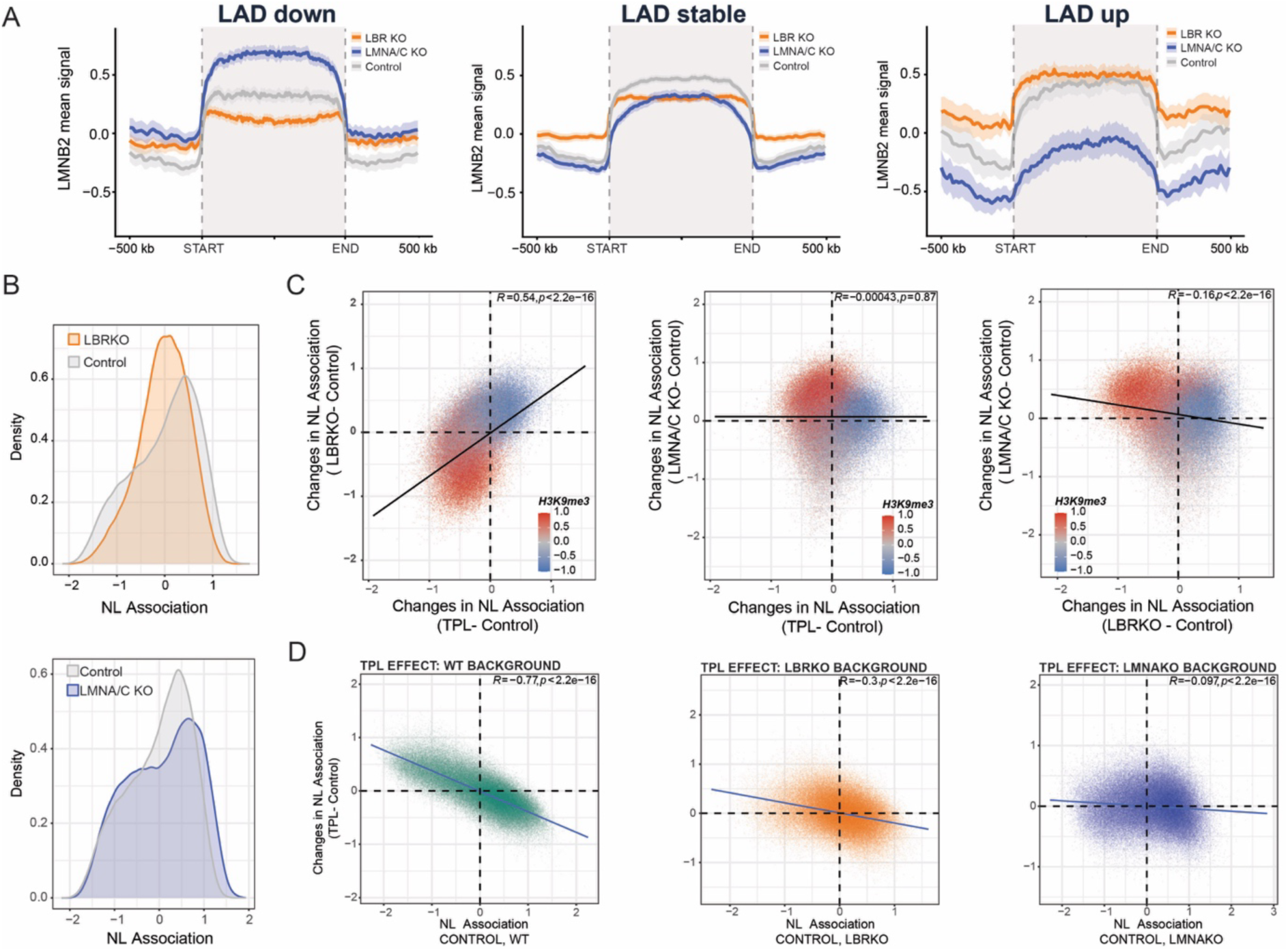
Opposite roles of LBR and LMNA/C on LAD/iLAD partitioning. A) Metaplot showing average pA-DamID signal for LMNB2 at LADs responsive or not to transcription inhibition and identified in Figure 4A for control (grey), LMNA/C KO (blue), and LBR KO (orange) RPE1 cells. LADs were length-normalized, and the average antibody signal was computed. The start and end of the normalized LADs are shown by dashed lines, and LAD regions are colored in grey. 500 Kb of not-normalized iLAD regions upstream and downstream of iLADs are shown in white. The shaded area around the average antibody signal represents 95% confidence intervals. Results are from 2 biological replicates in RPE1 cells. B) LMNB2 signal distribution for control (grey) and LBR KO (orange, *top panel*) and LMNA KO (blue, *bottom panel*). Results are from 2 independent biological replicates. C) Correlation scatter plots of 20 kb genomic bins. *Left panels*: differential LMNB2 score (TPL - control, x-axis) and differential LMNB2 score for LMNA/C KO (LBR KO - control, y-axis). *Middle panels*: differential LMNB2 score (TPL - control, x-axis) and differential LMNB2 score for LMNA/C KO (LBR KO - control, y-axis). *Right panels*: differential LMNB2 score (LBR KO - control, x-axis) and differential LMNB2 score for LMNA/C KO (LBR KO - control, y-axis). *Top panels:* bins are colored by H3K9me3 levels in control cells. D) Correlation scatter plots of 20 kb genomic bins for LMNB2 score in control cells (x-axis) and differential LMNB2 score (transcription inhibitor - control, y-axis) for TPL treatment in WT (green, three replicates), LBR KO (orange, two replicates), and LMNA KO (blue, two replicates) cells. The blue line represents a linear model; the Pearson correlation and *p*-value are shown in the plots.

#### LMNA/C and LBR differentially regulate genome-wide LAD/iLAD partitioning

To further investigate the interplay between transcription and lamina tethering mechanisms at the genome-wide level, we compared the effects of LMNA/C and LBR loss on chromatin-NL interactions at a global level. Similar to transcription inhibition (**Figure 1B**), LBR knockout induced a shift in LMNB2 signal distribution from a bimodal to a more unimodal profile, indicating reduced compartmentalization between LADs and iLADs (**Figure 6B**).

In contrast, LMNA/C knockout slightly increased the segregation of these domains, resulting in a more pronounced partitioning between LADs and iLADs (**Figure 6B**). Consistent with these observations, changes in chromatin-NL interactions following LBR loss positively correlate with those induced by transcription inhibition, whereas LMNA/C knockout shows a tendency towards anti-correlation, particularly in H3K9me3-enriched regions (**Figure 6C**).

Together, these findings demonstrate that transcription is a key determinant of chromatin–NL interactions and that its effects on genome positioning are mediated through distinct and opposing tethering mechanisms involving LBR and LMNA/C.

### Modulation of chromatin-NL contacts by Transcription requires LMNA and LBR

#### Partial Epistatic effects of LBR KO and transcription inhibition

To determine whether transcription can modulate chromatin-NL contacts via the two tethering mechanisms, we treated LBR KO cells and LMNA KO cells with TPL. Both cell lines showed TPL-induced degradation of RNA polymerase II (PolII) comparable to that in WT cells, suggesting that the drug is equally effective across the three cell lines (**Supplementary Figure 4D**). Despite similar PolII degradation triggered by TPL, the effects of transcriptional shutdown on chromatin-NL contacts in the LBR KO background were consistently milder, as revealed by a flattening of the slope of the correlation between NL association in untreated cells and changes induced by transcription inhibition, compared to the WT background (**Figure 6D**) and also by a reduction in the spread of the changes induced by the inhibitor (**Supplementary Figure 5G**). Given the strong correlation between changes induced by TPL and those induced by LBR KO (**Figure 6C**), these data indicate that loss of LBR-mediated tethering is partially epistatic to TPL-induced effects.

#### LMNA is required for Transcription modulation of chromatin-NL contacts

Strikingly, when we performed a similar analysis in LMNA KO cells, we found that the TPL effect was even further reduced, with a nearly horizontal slope in the correlation between NL association in untreated cells and changes induced by transcription inhibition (**Figure 6D**) and with the most reduced spread of the changes induced by the inhibitor across the three genetic backgrounds (**Supplementary Figure 5G**). Thus, the ability of transcription to shape chromatin-NL contacts appears to rely heavily on LMNA-mediated anchoring, and in its absence, chromatin-NL interactions become less sensitive to transcription shutdown.

Thus, the ability of transcription to control and shape chromatin-NL interactions depends on the presence of both tethers, with LMNA playing a more prominent role.

## DISCUSSION

The relationship between transcription and genome positioning at the nuclear lamina has remained conceptually elusive. Our results provide additional evidence that transcription is a major regulator of genome–NL interactions and shed light on the dynamics and mechanistic determinants of genome-NL interactions.

### High plasticity of chromatin-NL contacts

We show that acute transcription shutdown induces rapid and reversible rewiring of chromatin–NL contacts. Genome–lamina interactions are remodeled within one hour of transcription inhibition and largely restored after wash-off. This large-scale spatial repositioning occurs with minimal changes in heterocromatin state. This supports a model, consistent with previous works ^15,40,41^, in which spatial genome reorganization can precede, or be partly uncoupled from, chromatin-state remodeling. Thus, transcription is one of the key forces continuously maintaining LAD/iLAD partitioning, and lamina attachment appears to be an actively maintained state that can be rapidly released or reinforced before major chromatin-state changes become evident.

### Transcription as an antagonist of NL association

Previous work showed that gene activation within LADs can detach the corresponding transcription unit together with a limited surrounding region ^21^. Conversely, truncation of gene transcription by insertion of a premature stop codon increases NL association ^21^. Here, we observe a similar phenomenon at broader scale: acute transcription inhibition promotes reattachment of highly active genes and small neighboring iLADs, particularly near stably anchored LADs. These results support the idea that transcription acts as a local antagonist of lamina association, as previously proposed ^1,19–21,34^, while also showing that this effect is strongly constrained by genomic context. Proximity to a pre-existing LAD border and baseline NL affinity increase the likelihood of reattachment, suggesting that transcription-sensitive repositioning is facilitated by nearby stable anchors rather than occurring uniformly across the genome. These findings are consistent with a “zipper-like” model of LAD extension based on multiple weak and partly redundant interactions distributed across domains ^6,28,41^.

At single-gene and single-LAD levels, perturbing chromatin–NL contacts often induces compensatory movement of surrounding regions ^21,41^. However, our data show that, upon global transcription inhibition, gains and losses of NL association do not involve the same genomic regions or immediately adjacent regions. LAD weakening does not occur in LADs undergoing extension, indicating that these events are likely distinct and controlled by different mechanisms. One possible explanation for why H3K9me3-enriched LADs are displaced from the lamina upon transcription inhibition is limited space at the nuclear periphery. Previous single-cell DamID data suggested that the lamina may not have sufficient capacity to accommodate all genome simultaneously ^6^. Thus, when transcription inhibition causes many new regions to gain affinity for the NL, H3K9me3/LBR-associated LADs may be preferentially displaced as a consequence of competition for limited peripheral space.

### Different sensitivities of LMNA and LBR-tethered regions

A major conceptual advance from this work is that transcription-sensitive genome–lamina interactions reflect two distinct tethering pathways. LBR and lamin A/C are separable heterochromatin-tethering systems that can act sequentially during development ^27,42^. Our data show that LBR- and LMNA/C-mediated tethering differ in their acute responses to transcriptional perturbation within the same cellular context. Regions that lose NL association upon transcription shutdown are preferentially sensitive to LBR loss, whereas regions gaining NL contacts, particularly small iLADs near stable NL anchoring, are associated with LMNA/C-tethered regions. Thus, LBR and LMNA/C do not simply provide redundant anchoring capacity, but define mechanistically distinct tethers with different transcriptional dependencies. Active transcription appears to oppose incorporation of euchromatic inter-LAD regions into LMNA/C-associated peripheral domains, whereas LBR-dependent heterochromatin requires a transcription-competent nuclear environment to maintain stable lamina association.

### A possible mechanism involving LBR and LMNA

We propose that transcription continuously counterbalances LMNA/C-dependent peripheral capture of active chromatin while, directly or indirectly, sustaining LBR-dependent tethering of H3K9me3-marked LADs. In this scenario, LMNA/C may act as a barrier or buffer of transcriptional forces, whereas LBR tethering may depend on them for optimal binding. Transcription may therefore contribute differently to genome organization across cell types or differentiation stages, depending on whether they rely more on LMNA/C or LBR for heterochromatin tethering.

We show that both LMNA and LBR are required for a strong transcription-inhibition effect on chromatin–NL contacts. This may occur for two different reasons. For LBR, loss of the tether mimics transcription shutdown by triggering a similar de-partitioning effect, causing a partial epistatic phenotype. For LMNA, absence of this tether may lock the genome in a transcription- insensitive state. Loss of LMNA-mediated boundaries at highly transcribed iLADs may deprive these regions of stable anchoring that favors the “zipper-like” mechanism of LAD extension. At the same time, in LBR-dependent regions, LMNA may compete with or buffer LBR-mediated tethering; in its absence, LBR may bind too strongly and lose transcription sensitivity.

Several mechanistic questions remain open. We do not yet know whether transcription affects tethering through polymerase occupancy, DNA supercoiling, chromatin condensation, or transcription-coupled regulators that modulate LMNA/C- or LBR-based anchoring. Likewise, the basis by which transcription inhibition destabilizes LBR-dependent LADs while favoring LMNA/C-proximal iLAD absorption remains unclear. Whether and how LBR binds more strongly in the absence of LMNA/C will also require further investigation. Finally, a subset of LADs appears insensitive to transcription inhibition and to LMNA/C and LBR knock-out, suggesting that other tethers ^39^ or redundancy between LMNA and LBR are likely involved.

In summary, our study identifies transcription as a central regulator of genome organization at the nuclear lamina and shows that its effects are mediated by two distinct, opposing tethering pathways. While this manuscript was under preparation, a study showing that balancing transcription and cohesin activities regulates the positioning of genes proximal to LAD borders was published ^34^. Together, these and our data support a model in which LAD/iLAD partitioning is not fixed, but continuously shaped by the balance between transcription, loop extrusion, differential LMNA/C- and LBR-mediated heterochromatin tethering, highlighting the complexity of genome association with the NL.

## MATERIALS AND METHODS

### Cell lines and cell culturing

Two hTERT-immortalized RPE1 cell lines were used in this study. An RPE1 cell line expressing the SunTag system ^21,43^ was used for TPL and FLV treatments, as well as for mapping chromatin-NL contacts and heterochromatin components. This cell line was maintained in DMEM (Gibco), 10% FBS and 1% penicillin-streptomycin. A second RPE1 cell line expressing inducible Cas9 ^44^ was used to generate LBR and LMNA knockout cells and for all other experiments. This cell line was cultured in DMEM:F12 (Gibco, 11330032), 10% FBS and 1% penicillin-streptomycin. All cell lines were routinely tested for Mycoplasma contamination.

### Transcription inhibition

Triptolide (Merck-Sigma, T3652) and Flavopiridol (Merck-Sigma, F3055) were dissolved in DMSO at a concentration of 10 mM, aliquoted and stored at -80°C. Cells were treated with 2 μM of each drug for the indicated times. Drug-equivalent volumes of DMSO were used as control, corresponding to a final concentration of 0.02%.

### Antibodies used

The following antibodies were used: LMNB2 (Abcam, ab8983), LMNB1 (Abcam, ab16048), LMNA/C (SantaCruz Biotechnology, sc-376248), LBR (Abcam, ab32535), H3K9me3 (Diagenode, C15410193), H3K9me2 (Active Motif, 39239), H3K27me3 (Diagenode, C15410195), RPB1 (RNA Polymerase II, Santacruz Biotechnology, sc-11798).

### Generation of LBR and LMNA knock-out in RPE1 iCUT

LBR and LMNA knock-out RPE1 cell lines were generated as previously described ^38^.

### pA-DamID

pA-DamID maps were generated as previously described ^29^. Briefly, one million RPE1 cells were harvested by centrifugation at 500 g for 3 min and sequentially washed with ice-cold PBS and digitonin wash buffer (D-Wash) composed of 20 mM HEPES-KOH pH 7.5, 150 mM NaCl, 0.5 mM spermidine, 0.02% digitonin and Complete Protease Inhibitor Cocktail. Cells were incubated for 2 hours at 4°C under rotation in 200 μL D-Wash containing the antibody at a 1:200 dilution, followed by one wash with D-Wash. When the primary antibody was not raised in rabbit, cells were incubated with Rabbit-anti-mouse IgG diluted 1:200 in D-Wash buffer (Abcam, ab6709), followed by a wash step. The incubation was then repeated using a 1:200 pA-Dam solution, corresponding to approximately 60 Dam units as determined by calibration against Dam enzyme from NEB (#M0222L), followed by two washes. Dam activity was induced by incubating cells for 30 min at 37°C in 100 μL D-Wash supplemented with 80 µM SAM as methyl donor, under gentle shaking at 500 rpm.

For each condition, an additional one million cells were processed in D-Wash only and, during Dam activation, incubated with 4 units of Dam enzyme (NEB, M0222L). Dam-only control samples were used to normalize for DNA accessibility and amplification biases, as described ^45^. Genomic DNA was isolated and processed as for DamID, except that DpnII digestion was omitted. Libraries were sequenced either as 65-bp reads on a HiSeq 2500 or as 100-bp reads on a Novaseq platform.

For HiSeq libraries, library preparation was carried out as previously described ^29^. For Novaseq libraries, the following modified protocol was used. Genomic DNA was isolated using Bioline, BIO-52067, and approximately 500 ng were digested with DpnI (10 U, NEB, R0176L) in 1X CutSmart Buffer for 8 h at 37°C, followed by 20 min at 80°C, in a total volume of 10 µL. A-tailing was performed by adding 5 µL of A-tailing mix, consisting of 0.5 µL Cutsmart buffer 10X, 0.25 µL Klenow 50 U/µL (NEB, M0212M), 0.05 µL dATP 100 mM and 4.2 µL H2O, and incubating for 30 min at 37°C followed by 20 min at 75°C. Adapters were ligated by adding 15 µL ligation mix, containing 3 µL T4 Ligase Buffer 10X, 0.5 µL T4 ligase (5 U/µL, Roche, 10799009001), 0.25 µL x-Gene Stubby Adapter 50 µM (IDT) and 11.25 µL H2O. Samples were incubated for 16 h at 16°C and then for 10 min at 65°C. The ligation reaction was purified using 1.6 volumes of a custom-made solution containing SpeedBead Magnetic Carboxylate beads (Cytiva, 65152105050350), 18% PEG8000, 10 mM Tris-Cl pH 8.0, 1 mM EDTA, 1 M NaCl and 0.05% Tween. Finally, methyl-indexed PCR was performed by mixing 4 µL purified ligated DNA with x-Gen Dual combinatorial Indexes (IDT), at a final concentration of 125 nM, and MyTaq RedMix (Bioline, BIO-25048), in a final volume of 40 µL. The PCR program was as follows: 1 cycle of 1 min at 94°C; 14 cycles of 30 sec at 94°C, 30 sec at 58°C and 30 sec at 72°C; and 1 cycle of 2 min at 72°C.

Calibrated pA-DamID was performed as previously described ^32^. RPE1 cells were spiked with 20% mouse embryonic fibroblasts before proceeding with pA-DamID. In this experiment, LMNB1 pA-DamID was performed because this antibody worked efficiently in mouse cells, unlike the LMNB2 antibody.

### Bioinformatic analysis of pA-DamID data

pA-DamID data were processed as described ^14^. Briefly, adapter sequences were removed using cutadapt 1.11, and the remaining genomic DNA sequences were aligned to hg38 using bwa mem 0.7.17. Subsequent steps were performed using custom R scripts. Reads with a mapping quality of at least 10 and overlapping the ends of a GATC fragment were counted in 20-kb genomic bins. Counts were normalized to 1 million reads per sample, and a log2-ratio over the Dam control sample was calculated using a pseudo count of 1. At least two biological replicates were generated for each experimental condition, and the average score was used for downstream analyses.

#### Differential LAD calling

Differential LAD calling was performed as previously described ^29,32^. Briefly, to identify differential LADs, log2(LMNB2 : Dam) data were converted to z-scores, with a mean of zero and a standard deviation of one, to correct for differences in the dynamic range of pA-DamID signals between experiments while maintaining identical data distributions. LADs were identified in the control sample using a hidden Markov model applied to the average NL interaction profile between biological replicates (https://github.com/gui11aume/HMMt). The LAD score was calculated as the mean signal of the z-scaled data tracks. Significant changes were identified using a modified limma-voom approach ^46^. Significance was defined as a Benjamini-Hochberg adjusted P-value below 0.05.

#### Analysis at LAD borders

LADs defined as described in the section “Differential LAD calling” were used to visualize the consensus NL interaction profile around LAD borders. Average signal at LAD borders was calculated using 20-kb binned genomic tracks from pA-DamID (Log2(Ratio)). LAD flanks were extracted from the LAD models and used as LAD borders. LAD flanks located within 50 kb of chromosome ends were excluded, as these correspond to chromosome ends rather than actual LAD borders. Next, the closest LAD border was determined for each 20-kb genomic bin. The relative position from the closest LAD border was then used to calculate the mean score and the 95% confidence interval of the mean. Using the closest LAD border for each genomic bin ensured that each position was considered only once, including positions located near multiple LAD borders. LADs and iLADs smaller than 50 kb were excluded from this analysis because they contain only two normalized 20-kb bins and would therefore contribute only a single data point on one side of the LAD border, likely increasing noise in the resulting consensus profile.

#### Genomic metaprofile analysis of LAD and iLAD

Genomic metaprofiles across LADs and iLADs were generated using 20-kb binned genomic signal tracks containing chromosome, start and end coordinates, together with the signal values for each condition. LAD and iLAD coordinates were provided as BED files and included domains classified as increased, decreased or stable after TPL or FLV treatment. Signal files were converted to GRanges objects, and genomic signal was extracted over each LAD or iLAD together with 500 kb upstream and downstream flanking regions. Coordinates extending below position 1 were adjusted to start at position 1.

Because LADs and iLADs differ in size, domains were length-normalized before averaging. For each domain, the upstream flank, LAD or iLAD body, and downstream flank were treated separately. The LAD or iLAD body was interpolated to 100 bins, whereas each 500-kb flanking region was interpolated to 50 bins. Signal normalization across bins was performed by linear interpolation; when only one bin was available, the signal value was repeated across the normalized bins. After normalization, all domains were aligned relative to their normalized positions, with upstream regions, domain bodies and downstream regions represented on a common scale.

For each normalized position and condition, the mean signal was calculated across all LADs or iLADs, together with the standard deviation, standard error and 95% confidence interval. Confidence intervals were calculated from the standard error using a normal approximation.

#### Analysis of calibrated pA-DamID data

Calibrated pA-DamID data were processed similarly to uncalibrated data, with the exception that reads were aligned simultaneously to the human hg38 and mouse mm10 genomes, and genomic bins of 250 kb were used instead of 20 kb. The larger bin size was required because the number of mouse reads was limited and larger bins provided more stable scaling estimates. Filtering based on a mapping quality of 10 ensured that only uniquely aligned reads mapping to either the human or mouse genome were retained for downstream analyses. After calculating the Log2(Ratio) over the Dam control using the combined genome reference, scaling factors were calculated to convert the mouse log2(ratios) to z-scores, as described in the section “different LAD scaling”. All mouse spike-ins originated from the same biological sample, and this scaling procedure ensured that all mouse profiles were normalized to the same dynamic range. The same scaling factors were then applied to the human log2(ratios) for each sample. In contrast to uncalibrated data, this approach allowed quantitative differences in dynamic range between experiments, caused by increased or decreased protein-DNA interactions, to be detected.

### PCA and Linear Regression Modelling

Genome-wide signal tracks were imported from bigWig files using pyBigWig and aggregated over 20-kb genomic bins across human chromosomes chr1–chr22, chrX and chrY. Control heterochromatin marks were loaded from pADamID profiles for H3K9me2, H3K9me3 and H3K27me3, and differential lamina association changes were loaded from the Triptolide and Flavopiridol contrast tracks. RNA-seq replicate tracks were also binned at 20 kb before merging. All tracks were merged into a single locus-level table.

For the multivariate analyses, control histone signals were represented as H3K9me2, H3K9me3 and H3K27me3 taken from the control profiles, while transcription was represented by the sum of the two RNA-seq replicates and log-transformed as log(RNAseq_1 + RNAseq_2 + 1). Missing values were filtered and remaining gaps were imputed as zeros for PCA and regression.

PCA was carried out by singular value decomposition on the 4-dimensional matrix of H3K9me2, H3K9me3, H3K27me3 and RNAseq. Score plots were generated for PC1–PC2 and PC1–PC3, with feature loadings projected as vectors to visualize the relative contribution of transcription and heterochromatin marks to the major axes of chromatin variation. One PCA visualization was colored by local point density (Figure 2 A top), or by the TPL-induced lamina association change (Figure 2 A bottom).

Least-squares regression was then used to predict treatment-induced NL contact changes (TPL and FLV) from the same four predictors. Ordinary least squares coefficients were estimated from the normal equations including an intercept, and coefficient uncertainty was computed from the residual variance and the inverse of X^T^ X. Model fit was assessed with R^2^ and Spearman rank correlation between observed and predicted values.

### Calibrated RNA-seq

RPE1 cells were treated with TPL for 3 hours, collected on ice by scraping, and spiked with 10% Drosophila S2 cells. Cell pellets were snap-frozen in liquid nitrogen and stored at -80°C. RNA was extracted using the NZY Total RNA Isolation Kit (Nzytek, MB13402) and stored at -80°C. RNA samples were submitted to Novogene for quality control, library preparation and sequencing. Libraries were prepared using the Novogene NGS RNA Library Prep Set (PT042). Briefly, messenger RNA was purified from total RNA using poly-T oligo-attached magnetic beads followed by ethanol precipitation. The precipitated RNA was mechanically fragmented by sonication using Covaris (Massachusetts, United States), and first-strand cDNA synthesis was performed using random hexamer primers, followed by second-strand cDNA synthesis. cDNA ends were converted to blunt ends using enzymes with exonuclease/polymerase activity, followed by adenylation of the 3′ ends and adapter ligation.

Size selection was performed to obtain a 250–300 bp insert strand-specific library, and adapter-ligated cDNA fragments were selectively enriched by PCR. After PCR amplification, products were purified using the AMPure XP system (Beckman Coulter, Beverly, USA). Sequencing was performed by Novogene on an Illumina NovaSeq X Plus platform (Illumina, USA) using a paired-end 150 bp sequencing strategy on a 25B lane. Approximately 40 million reads were generated for each condition. Three independent biological replicates were produced. RNA-seq data analysis.

Count tables were produced from raw reads with RSEM (v1.3.3), using Bowtie2 (v2.5.4) for read mapping against a combined transcriptome reference generated from the hg38 and dm6 genome assemblies and corresponding RefSeq gene annotations as downloaded from the UCSC Genome Browser download section. Chromosome, gene, and transcript identifiers were prefixed with species-specific labels to allow unambiguous assignment of reads. Differential expression analysis of human genes was performed using DESeq2 (v1.52.0), with size factors estimated from Drosophila genes and a design formula including treatment condition and replicate identity (∼ replicate + condition).

For metaprofiles analysis each library was scaled by a spike-in size factor defined as *s = R/d*, where *d* is its Drosophila read count and *R* the geometric mean of *d* across all libraries (MAPQ ≥ 30, excluding reads mapped with SAM flag 2820). Human-only coverage tracks were generated with deepTools (v3.5.6) bamCoverage using 50-bp bins, *--normalizeUsing None* and the corresponding *--scaleFactor.* Replicate tracks were averaged into per-condition mean tracks, which were then combined into log2 ratio tracks (T3- and T6-vs-CTRL) with pseudocount 0.01. Strand-oriented 1-bp boundaries were defined at LAD/iLAD interval starts and ends, and computeMatrix (reference-point mode, ±500 kb, 5-kb bins) was used to build orientation-harmonised coverage matrices for the six boundary categories. Metaprofiles were computed as the across-region mean per bin, with a 68% (≈ ±1 SEM) region-bootstrap band.

### Statistical analyses

Statistical analyses, when performed, are indicated in the text or figure legends.

## ACKNOWLEDGMENTS

We thank the NKI Genomics, Research High-Performance Computing, and Robotics & Screening core facilities; the DBS Flow Cytometry, Protein Expression and Purification, and Microbiology core facilities at Unimi; and the Genomic National Platform at Human Technopole for technical support. We thank members of our laboratories for stimulating discussions.

This work was supported by the European Union (ERC, GoCADiSC, 694466 to B.v.S.; AIRC-MSCA iCARE2 fellowship 800924, MSCA-IF 838555, Next generation EU-MUR MSCA Young Researcher, University of Milan PSR Linea 4 and PSR Linea 8, Human Technopole National Facilities ID2166943 to S.G.M; R.M. is supported by NWO Zwaartekracht (58588); M.M.F is supported by AIRC IG30264; G.P. and F.Z. were supported by Italian National Recovery and Resilience Plan (PNRR) project ELIXIRNextGenIT (R0000010). Views and opinions expressed are those of the authors only and do not necessarily reflect those of the European Union or the European Research Council. Neither the European Union nor the granting authority can be held responsible for them. Research at the Netherlands Cancer Institute is supported by institutional grants of the Dutch Cancer Society and the Dutch Ministry of Health, Welfare and Sport. The Oncode Institute is partially funded by the Dutch Cancer Society.

## AUTHOR CONTRIBUTIONS

Conceptualization: S.G.M, B.v.S. Methodology: M.B, S.G.M, A.G.M. Investigation: S.G.M., M.B. Formal Analysis: S.G.M, M.B, J.B, T.v.S, F.Z, G.P. Software: T.v.S. Data curation: S.G.M. Supervision: S.G.M and B.v.S. Writing - original draft: S.G.M with input from the others. Funding acquisition: S.G.M, B.v.S, R.M, MMF, G.P, F.Z. All authors have read and approved the final manuscript.

## SUPPLEMENTARY FIGURES

**Supplementary Figure 1.**
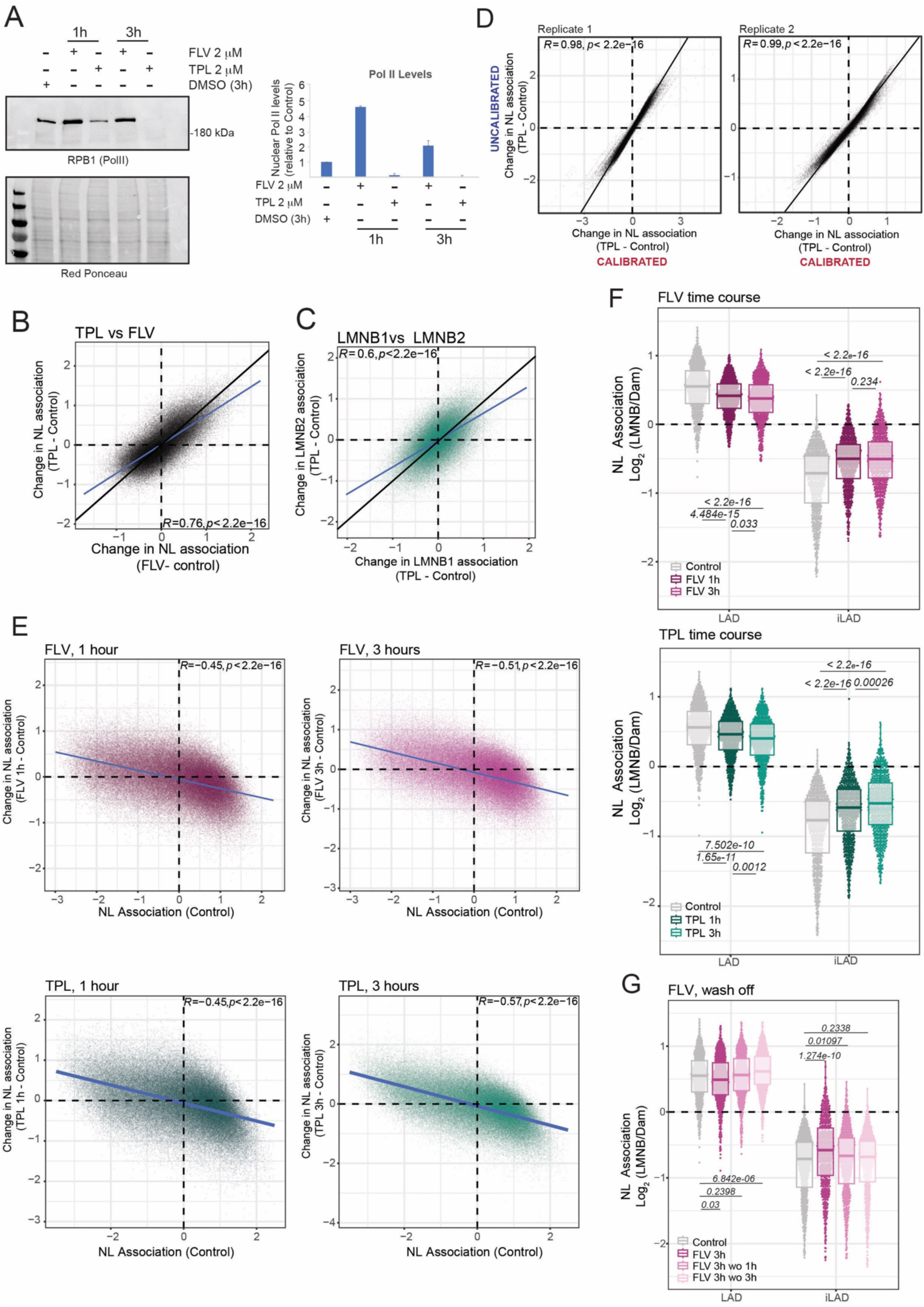
Transcription controls genome partitioning in LADs and iLADs. A) *Left:* Representative western blot analysis showing nuclear levels of RPB1, the largest subunit of RNA Polymerase II following TPL and FLV treatments performed at 1 and 3 hours, at 2 µM. *Right:* Quantification of RPB1 levels. Red ponceau staining was used to normalize for protein quantity. Data are normalized on control levels. Error bars represent standard deviation. Results are from two biological replicates. B) Correlation scatter plots of 20 kb genomic bins for differential LMNB2 (transcription inhibitor - control) for FLV-treated (x-axis) and TPL-treated (y-axis) cells. Results are from three biological replicates. C) Correlation scatter plots of 20 kb genomic bins for differential LMNB1 and LMNB2 scores (transcription inhibitor - control) for TPL-treated cells. Results are from two biological replicates. D) Genome-wide correlation of differential LMNB1 score (TPL treatment-control) for calibrated (mouse reads normalization following MEF spike-in) and uncalibrated data for each replicate. E) Correlation scatter plots of 20 kb genomic bins for LMNB2 score in control cells (x-axis) and differential LMNB2 score (transcription inhibitor - control, y-axis) for FLV and TPL treatments at 1 and 3 hours of treatment. Results are from two biological replicates for FLV and one for TPL. F) LAD and iLAD score following FLV or TPL treatment at 1 and 3 hours using LMNB2 mapping data. Results are from two biological replicates. G) Same as F, but for a FLV wash-off experiment. Results are from two biological replicates. *For all scatter plots in the figure, the blue line represents a linear model, and the black line represents the diagonal. Pearson correlation and p-value are shown in the plots*.

**Supplementary Figure 2.**
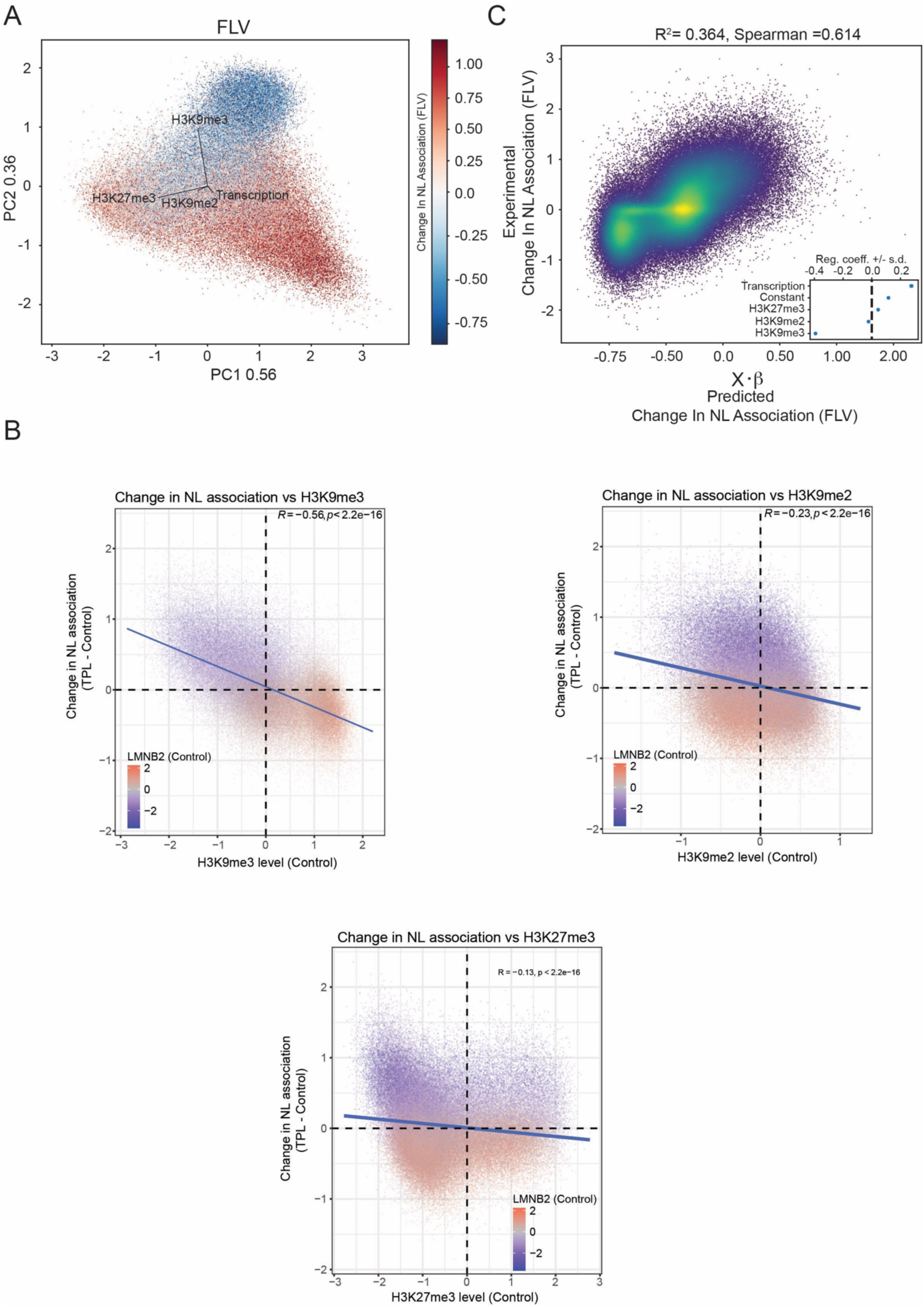
Chromatin composition is predictive of chromatin-NL changes. A) Top panel: PCA performed on 100-kb genomic bins using H3K9me2, H3K9me3 and H3K27me3 pA-DamID data together with RNA-seq data from RPE1 cells. H3K9me2, H3K9me3, H3K27me3 and log-transformed RNA-seq values were decomposed by singular value decomposition. PC1 and PC2 are shown and together explain 92% of the variability in chromatin composition. Dots are colored by changes in chromatin–NL interactions following FLV treatment relative to control conditions (FLV–Control; 3 h, 2 μM). Black arrows indicate the feature loadings for each marker. *Bottom panel:* Same PCA analysis as in A, with bins colored. B) Correlation scatter plot of 20-kb genomic bins for H3K9me3, H3K9me2, and H3K27me3 scores in control cells (x-axis) and differential LMNB2 score (TPL - control, y-axis). The blue line represents a linear model; Pearson correlation and *P* value are shown in the plot. C) Least-squares regression model of FLV-induced changes in chromatin–NL interactions. Observed changes in nuclear lamina association are plotted against values predicted by a linear model including an intercept plus H3K9me2, H3K9me3, H3K27me3 and RNA-seq. The model shows that transcription and H3K9me3-marked heterochromatin are predictive of gain and loss of chromatin–NL interactions following FLV treatment, respectively. R² and Spearman correlation are shown on the plot. The inset reports the estimated regression coefficients for each predictor, with ±2 standard-deviation intervals. Results are from two biological replicates.

**Supplementary Figure 3.**
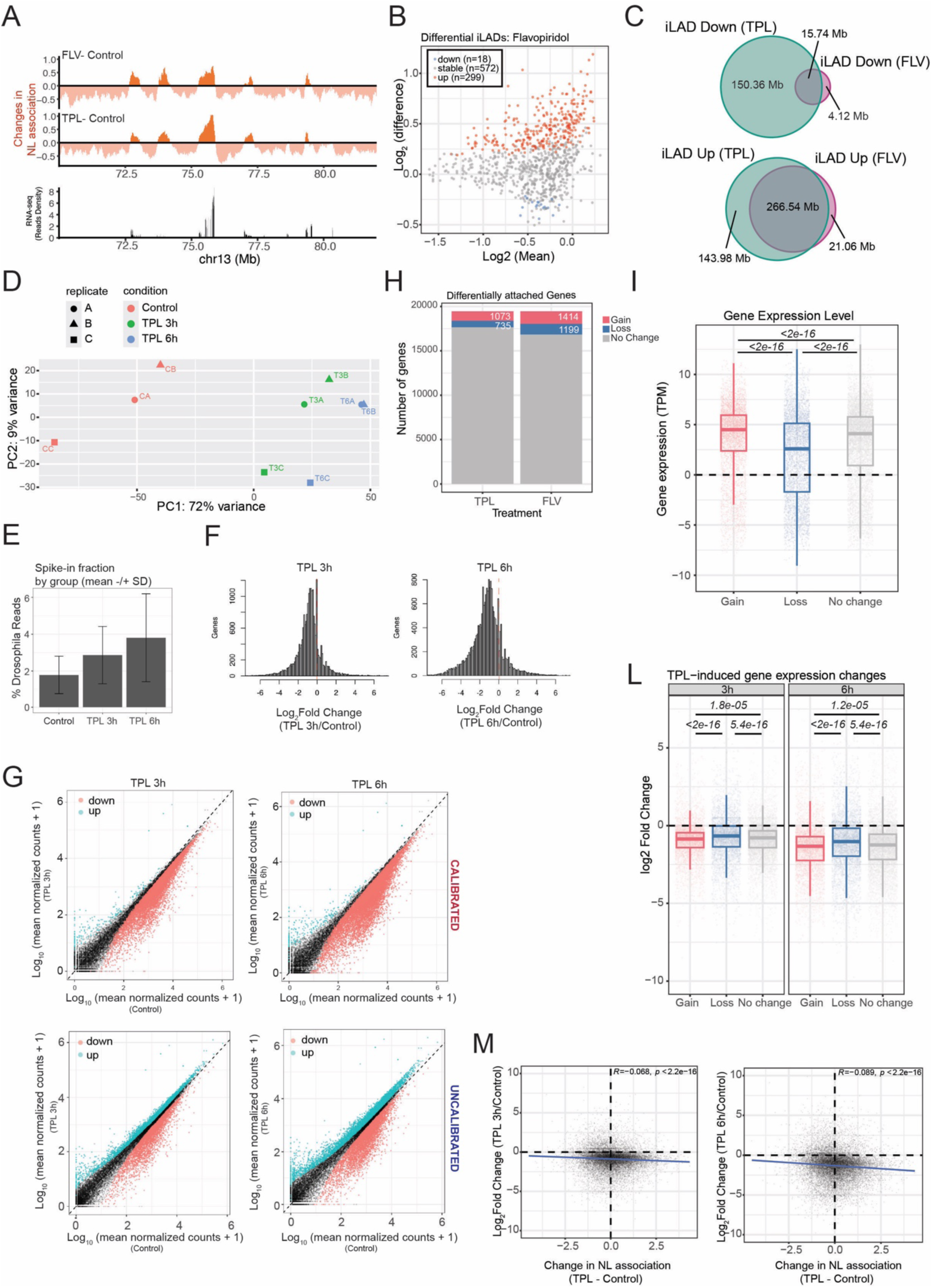
A) *Top panel*: Differential tracks (treatment - control, orange) of 12 Mb on chromosome 13, highlighting the gain and loss of genome-NL interactions following inhibition of transcription with the two different drugs. *Bottom panel*: RNA-seq tracks showing transcription level for genes in the same genomic area. B) Results of Voom-Limma analysis used to call iLADs that significantly gain or lose interaction with LMNB2 following FLV treatment. Results are from 4 biological replicates. C) Overlap in Mb for iLAD down and up following TPL (green) and FLV (purple) treatments. D) PCA analysis performed on RNA-seq reads from control cells and cells treated with TPL for 3 and 6 hours, and for the three different biological replicates. E) Bar plot showing the percentage of spiked-in Drosophila reads for the different experimental conditions. Error bars represent standard deviation. Results are from three biological replicates. F) Change in gene expression (expressed as Log2(Fold Change), TPL vs DMSO) for calibrated data at 3 and 6 hours. G) Genome-wide correlation of mean normalized counts between TPL treatment and control for calibrated (*top panels*) and uncalibrated data (*bottom panels*) for 3 hours (*left panels*) and 6 hours (*right panels*) of TPL treatment. H) Number of differentially attached genes following FLV and TPL treatments (3 hours, 2 μM). Gained and lost genes were defined by setting a cut-off of +/- 0.5 on the differential pA-DamID score (treatment - control). I) Gene expression levels, expressed in TPM, for genes that gain (red), lose (blue), or do not change (grey) chromatin-NL interactions following TPL treatment. L) Change in gene expression (expressed as Log2 Fold Change, TPL vs DMSO) for gain, loss, and no change genes at 3 (left panel) and 6 (right panel) hours of TPL treatment (2 μM). For H and I, the P values are based on Wilcoxon’s test. For I, P values are based on Wilcoxon’s test with the “less” hypothesis tested. M) Correlation scatter plot between changes in LMNB2 pA-DamID score (TPL - control; 3 hours in left panel, and 6 hours in right panel) and change in gene expression (expressed as Log2(Fold Change), TPL vs DMSO). The blue line represents a linear model; Pearson’s correlation and p- value are shown in the plots.

**Supplementary Figure 4.**
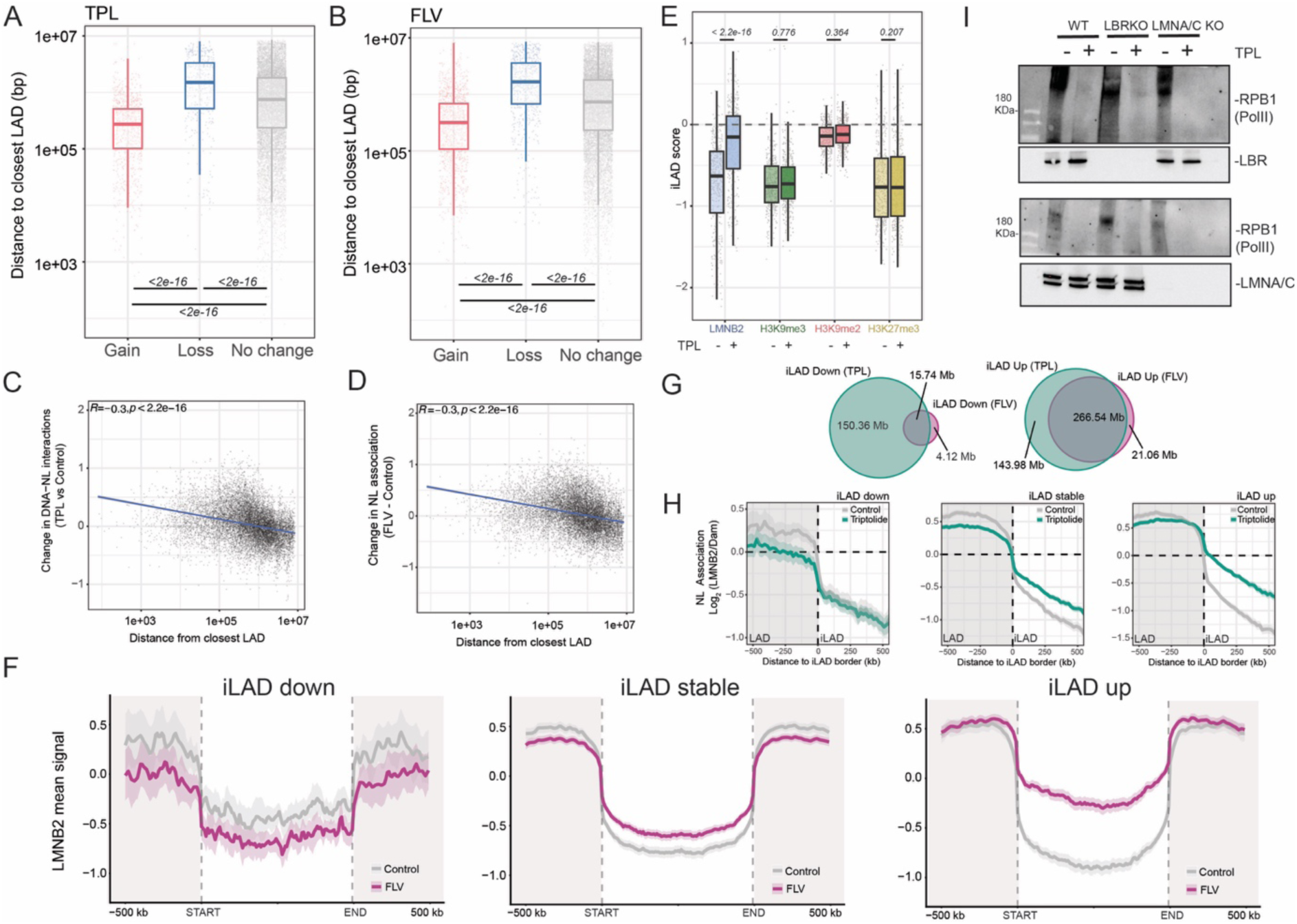
A) Distances from the closest LAD border, for genes that gain (red), lose (blue), or do not change (grey) chromatin-NL interactions following TPL treatment. LAD genes (Distance to LAD <=0) were excluded from this analysis. The P values are based on Wilcoxon’s test. B) Same as A, but for FLV treatment. C) Correlation scatter plots of genic distances from the closest LAD border in control cells (x-axis) and differential LMNB2 score (transcription inhibitor - control, y-axis) for TPL treatment (3 hours, 2 µM). The blue line represents a linear model; Pearson’s correlation and p-value are shown in the plots. D) Same as C, but for FLV treatment. E) LMNB2, H3K9me3, H3K9me2, and H3K27me3, average pA-DamID scores for control and TPL-treated cells for iLADs, showing a significant increase in interaction following TPL treatment. The P values are based on Wilcoxon’s test. F) Metaplot showing average pA-DamID signal for LMNB2 at iLADs up (*left*), stable (*middle*), or down (*right*), identified in Figure 3A, in control or FLV-treated cells (3 hours, 2 µM). Control is shown in grey, and FLV-treated samples are shown in purple. iLADs were length-normalized, and the average antibody signal was computed. The start and end of the normalized iLAD are shown by dashed lines. 500 kb of non-normalized LAD regions upstream and downstream of iLADs are shown in grey. The shaded area around the average antibody signal represents 95% confidence intervals. Results are from 4 biological replicates in RPE1 cells. G) Overlap in Mb for iLAD down and up following TPL (green) and FLV (purple) treatments. H) Average LMNB1 pA-DamID score around iLAD (down, stable or up) borders for control and TPL-treated samples. Genome-NL contact profiles are the average of 4 biological replicates. The solid line and the shaded area represent the mean signal and 95% confidence interval of the mean, respectively. I) Western blot analysis showing total levels of RPB1, the largest subunit of RNA Polymerase II, following TPL treatments performed at 3 hours, at 2 µM in RPE1 WT, LBR KO, and LMNA/C KO cells.

**Supplementary Figure 5.**
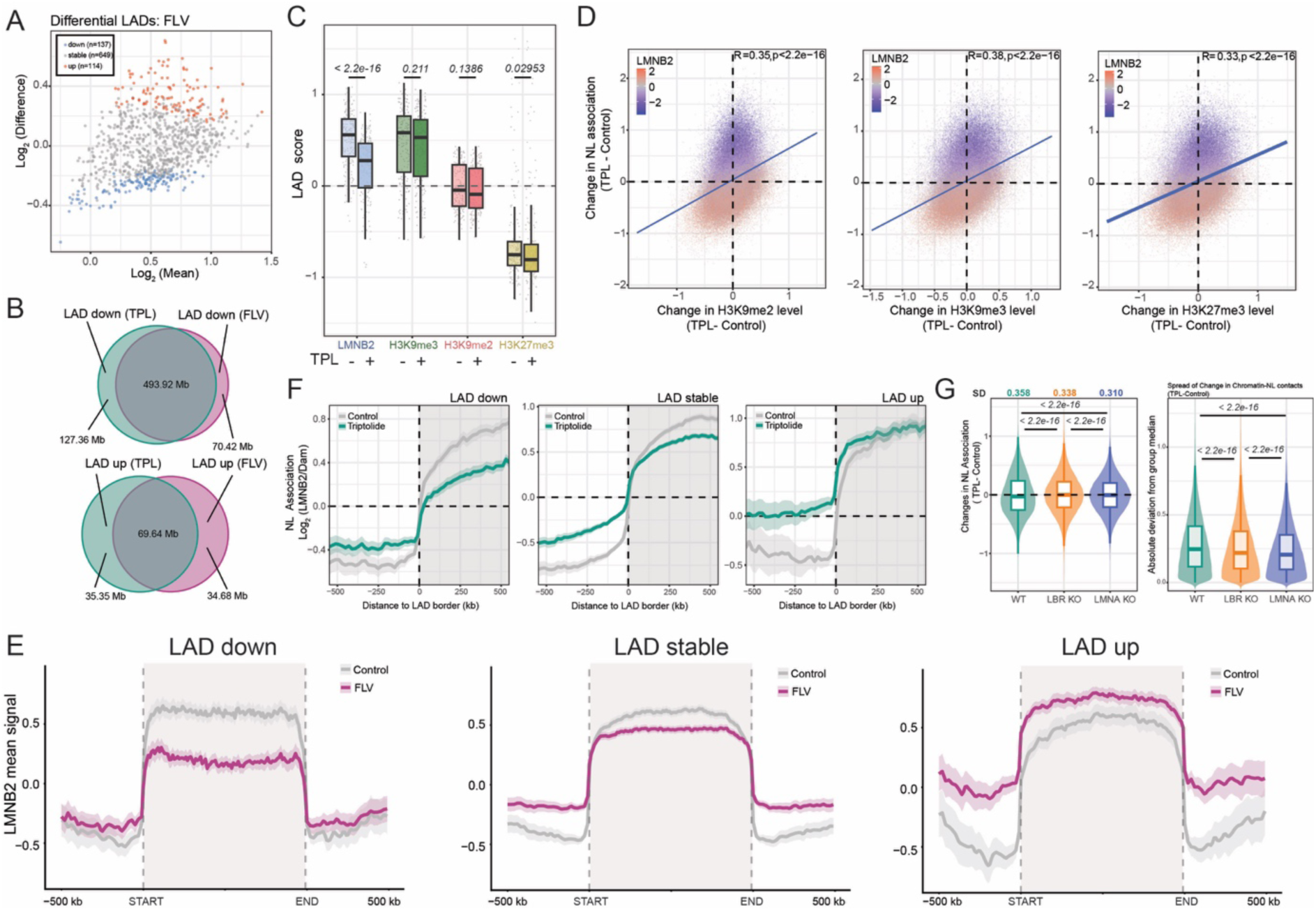
A) Results of Voom-Limma analysis used to call LADs that significantly gain or lose interaction with LMNB2 following FLV treatment (3 hours, 2 μM). Results are from 4 biological replicates. B) Overlap in Mb for LAD down and up following TPL (green) and FLV (purple) treatments. C) LMNB2, H3K9me3, H3K9me2, and H3K27me3, average pA-DamID scores for control and TPL-treated cells for LADs showing a significant decrease in interaction following TPL treatment. The P values are based on Wilcoxon’s test. D) Correlation scatter plot of 20-kb genomic bins for differential H3K9me3, H3K9me2, and H3K27me3 scores (x-axis, TPL - control) and differential LMNB2 score (y-axis, TPL - control). The blue line represents a linear model; Pearson’s correlation and p-value are shown in the plots. E) Metaplot showing average pA-DamID signal for LMNB2 at LADs down (*left*), stable (*middle*), or up (*right*), identified in Figure 4A, in control or FLV-treated cells (3 hours, 2 μM). Control is shown in grey, and FLV-treated samples are shown in purple. LADs were length-normalized, and the average antibody signal was computed. The start and end of the normalized LAD are shown by dashed lines. 500 Kb of not-normalized iLAD regions upstream and downstream of LADs are shown in grey. The shaded area around the average antibody signal represents 95% confidence intervals. Results are from 4 biological replicates in RPE1 cells. F) Average LMNB2 pA-DamID score around LAD (down, stable or up) borders for control and TPL-treated samples. Genome-NL contact profiles are the average of 4 biological replicates. The solid line and the shaded area represent the mean signal and 95% confidence interval of the mean, respectively. G) *Left:* Distribution of changes in chromatin–NL contacts induced by TPL treatment (TPL − control) in WT, LBR KO, and LMNA KO cells. Differences in the dispersion of Delta score distributions were assessed using a median-centered Levene’s test. Standard deviation value for each distribution is reported on top of the panel. *Right:* Distribution of absolute deviations from the group median following TPL treatment across the three genetic backgrounds. Pairwise t-tests were performed on absolute deviations from the group median to assess differences in dispersion between groups.

